# Transcriptional remodeling by OTX2 directs specification and patterning of mammalian definitive endoderm

**DOI:** 10.1101/2024.05.30.596630

**Authors:** LS Ee, D Medina-Cano, CM Uyehara, C Schwarz, E Goetzler, E Salataj, A Polyzos, S Madhuranath, T Evans, AK Hadjantonakis, E Apostolou, T Vierbuchen, M Stadtfeld

## Abstract

- Mouse and human pluripotent cells with multipurpose degron alleles establish versatile platforms to dissect cell type-specific functions of the pleiotropic transcription factor OTX2
- OTX2 controls molecular programs required for anterior-posterior patterning of the developing gut
- OTX2 establishes and maintains chromatin accessibility at distinct distal gene regulatory elements in definitive endoderm
- OTX2 functions as a patterning factor across different germ layers and species

The molecular mechanisms that drive essential developmental patterning events in the mammalian embryo remain poorly understood. To generate a conceptual framework for gene regulatory processes during germ layer specification, we analyzed transcription factor (TF) expression kinetics around gastrulation and during in vitro differentiation. This approach identified Otx2 as a candidate regulator of definitive endoderm (DE), the precursor of all gut-derived tissues. Analysis of multipurpose degron alleles in gastruloid and directed differentiation models revealed that loss of OTX2 before or after DE specification alters the expression of core components and targets of specific cellular signaling pathways, perturbs adhesion and migration programs as well as de-represses regulators of other lineages, resulting in impaired foregut specification. Key targets of OTX2 are conserved in human DE. Mechanistically, OTX2 is required to establish chromatin accessibility at candidate enhancers, which regulate genes critical to establishing an anterior cell identity in the developing gut. Our results provide a working model for the progressive establishment of spatiotemporal cell identity by developmental TFs across germ layers and species, which may facilitate the generation of gut cell types for regenerative medicine applications.

## Introduction

Pluripotency exit and morphogenetic changes during gastrulation that result in the acquisition of the germ layers ectoderm, mesoderm, and endoderm are central processes for tissue specification in the post-implantation mammalian embryo and set the stage for later organogenesis (Bardot and Hadjantonakis, 2020; Tam and Behringer, 1997). Genetic studies in model organisms have identified numerous modulators of gastrulation, including regulators of signaling pathways that instruct germ layer identity (Morgani and Hadjantonakis, 2020) and cell surface molecules that promote correct morphogenetic cell migration and adhesion (Muncie et al., 2020). In addition, DNA-binding transcription factors (TFs) have been recognized as important regulators of germ layer and organ specification (Cui et al., 2018; Tam and Loebel, 2007) whose dysregulation can drive developmental and congenital disorders.

Recent single-cell profiling experiments have provided high-resolution atlases of transcriptional changes associated with gastrulation and the emergence of specific germ layers in mouse and human embryos (Nowotschin et al., 2019; Pijuan-Sala et al., 2019; Qiu et al., 2022; Tyser et al., 2021; Zeng et al., 2023). Yet, our understanding of how specific TFs coordinate the transcriptional remodeling required for germ layer specification and organ development at the molecular level remains superficial. This lack of knowledge is due to (i) the inaccessibility of the post-implantation mammalian embryo, (ii) the associated difficulty in generating sufficient cellular material for in-depth molecular characterization of TF functions, and (iii) limitations of existing gene knockout (KO) models concerning specificity and spatiotemporal control, suggesting a need for more tractable experimental systems.

Embryonic stem cells (ESCs) are attractive tools for studying developmental processes (Gaertner et al., 2019). Still, standard mouse ESC (mESC) differentiation protocols are highly variable and yield disorganized and/or heterogeneous cell populations that inadequately mirror embryonic patterning processes (Bernardo et al., 2011; Drukker et al., 2012; Yu et al., 2011). Human ESCs (hESCs) achieve more homogenous differentiation responses upon suitable stimulation (Hawkins et al., 2021; Loh et al., 2014; Pan et al., 2020), possibly because they resemble a post-implantation “primed” state of development (Tesar et al., 2007). Accordingly, converting naïve mESCs into a primed state allows near-homogenous, signaling-mediated differentiation into tissues such as definitive endoderm (DE), the precursor of all gut-derived organs (Medina-Cano et al., 2022). For the interrogation of protein function, in-frame fusions of degron tags such as FKBP12^F36V^ (referred to as the dTAG system) (Nabet et al., 2018) to proteins of interest has emerged as a powerful approach to enable acute, controlled and specific depletion of candidate factors (Jaeger and Winter, 2021; Prozzillo et al., 2020), which has been applied to address developmental questions (Abuhashem et al., 2022; Bisia et al., 2023). Integrating degron alleles into synchronized directed differentiation regimens, therefore, in principle, represents a tractable and versatile experimental platform to dissect the function of developmental TFs at high temporal resolution.

OTX2 is a conserved homeobox TF with pleiotropic functions during vertebrate development (Beby and Lamonerie, 2013). In mouse and other animal models, OTX2 is required for correct migration and inductive properties of the anterior visceral endoderm (Rhinn et al., 1998), an extra-embryonic tissue required for establishing anterior-posterior identity in the mammalian embryo. Subsequently, OTX2 by cell-autonomous and non-autonomous mechanisms (Rhinn et al., 1999) controls the specification and patterning of the neuroectoderm (Acampora et al., 1995) and later the formation of specific neuroectoderm derivatives (Beby and Lamonerie, 2013). Gastrulation-stage mouse embryos deficient for OTX2 manifest with defects in non-ectodermal tissues, including foregut and heart abnormalities (Ang et al., 1996; Jin et al., 2001; Matsuo et al., 1995) suggesting OTX2 functions in germ layers beyond ectoderm.

By mining published and unpublished genomics datasets, we identified evidence for a direct role of OTX2 in the formation and patterning of the mammalian DE and primary gut tube. Systematic assessment of embryonic endoderm formation from mouse and human primed pluripotent stem cells harboring “multipurpose” (degradation/immunoprecipitation/visualization) degron alleles showed that stage-specific depletion of OTX2 resulted in abnormal DE with impaired ability for anteriorization and foregut differentiation. Transcriptional functions of OTX2 in the DE include the activation of specific gut tube-associated downstream TFs, balanced expression of WNT and FGF/ERK signaling components, and repression of non-endodermal genes. Gene activation by OTX2 entails OTX2-dependent gain of chromatin accessibility at distal enhancer elements harboring canonical binding motifs, some of which are primed for activation by OTX2 binding in the epiblast. In contrast, OTX2-dependent gene repression proceeds via binding to low-affinity sites and by indirect mechanisms. Our results suggest that OTX2 is a central component of a transcriptional program for anteriorization during peri-gastrulation phases of development that is conserved across germ layers and species.

## Results

### Integrative analysis of in vivo and in vitro datasets identifies candidate regulators of germ layer specification

We analyzed available genomics and phenotypic data to identify candidate transcriptional regulators of germ layer specification among a comprehensive list of 1,682 murine TFs (Garipler et al., 2022). We set as criteria **I)** strong upregulation – fold change (FC)>5 and adjusted p-value (padj) <0.05 – during the transition of mESCs to epiblast-like stem cells (EpiLCs) which resemble the early post-implantation epiblast (Hayashi et al., 2011) (**Table S1**), consistent with a role in preparing for germ layer specification; **II)** sustained elevated expression throughout formation and patterning of at least one germ layer in the mouse embryo based on single-cell RNA sequencing (scRNA-seq) (Nowotschin et al., 2019; Pijuan-Sala et al., 2019) and **III)** developmental organ phenotypes in KO mice in the Mouse Genome Database (Bult et al., 2019). This approach resulted in the identification of a group of seven candidate TFs that met all criteria (**Fig. S1A**). Among these candidates, *Otx2* – encoding an essential and highly conserved homeobox TF (Ang et al., 1996; Matsuo et al., 1995) – demonstrated the most significant upregulation during the transition from naïve to primed pluripotency (**Table S1**). While the importance of OTX2 for development of the anterior visceral endoderm and anteriorization of the neuroectoderm (NE) have been documented (Acampora et al., 1995; Hever et al., 2006; Simeone, 1998), we also observed *Otx2* expression along the trajectory of early gut tube development in the mouse embryo (post-implantation epiblast ® anterior primitive streak ® definitive endoderm (DE) → anterior gut tube) (**Fig.1A,B**). *Otx2* levels in DE were comparable to those in VE and NE (**Fig. S1B**), consistent with an unexplored role of this TF in the embryonic endoderm.

**Figure 1.**
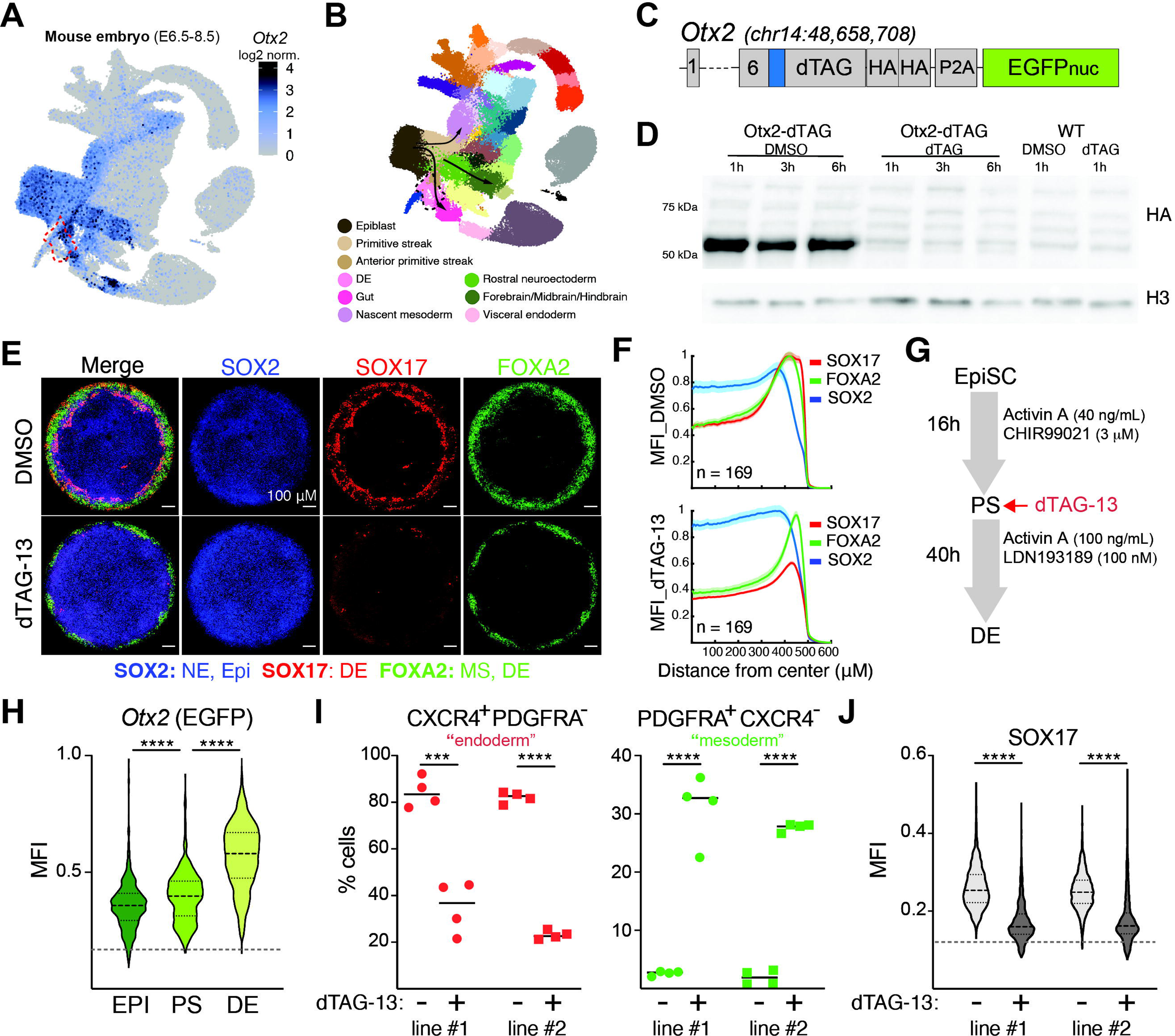
OTX2 depletion impairs DE development in murine models of germ layer specification. A. *Otx2* expression levels in gastrulation-stage (E6.5-8.5) mouse embryos (modified after Pijuan-Sala et a., 2019). The red dotted line indicates the position of DE. B. Tissue annotation of gastrulation-stage mouse embryos with tissues expressing *Otx2* listed. Arrows indicate different developmental trajectories along which *Otx2* is expressed. C. Design of the murine OTX2-dTAG allele. Coordinates indicate the position of the *Otx2* STOP codon (mm10), which was replaced with a multipurpose cassette. Blue box indicates a linker peptide. D. HA Western blotting (WB) after treatment of OTX2-dTAG EpiLC or EpiLCs derived from parental wildtype (WT) mESCs with dTAG-13 or DMSO for the indicated periods of time. Asterisk indicates a non-specific band. E. Representative IF images after staining anterior micropatterns formed in presence (top) or absence (bottom) of OTX2 for marker proteins of indicated lineages. F. Average marker intensity (+/- standard error of the mean) at different radial positions in anterior micropatterns. N = colonies analyzed. G. Protocol for directed differentiation of EpiSCs into DE, indicating compounds applied and timing of dTAG-13 administration for experiments shown in **Figs. 1I,J**. H. Quantification of Otx2-EGFP levels at indicated stages of directed differentiation. Dotted line indicates level of background fluorescence measured in WT cells. (****)p<0.0001 with one-way ANOVA with Tukey’s multiple comparison test. I. Quantification by flow cytometry of CXCR4^+^PDGFRA^-^ (DE) and PDGFRA^+^CXCR4^-^ (mesoderm) cells following directed differentiation in control (DMSO) or OTX2-depleted (dTAG-13) conditions. (***)p<0.001 and (****)p<0.0001 with unpaired t-test. N=4 separate cultures from each of two independent cell lines. J. Quantification of IF imaging of SOX17 (an endodermal marker) in DE derived from two independent OTX2-dTAG EpiSC lines cultured in absence or presence of dTAG-13. (****)p<0.0001 with unpaired t-test. N=4 separate cultures per cell line.

To enable studies into developmental functions of OTX2, we used Cas9-faciliated gene targeting to generate several mESC lines homozygous for a “multipurpose” degron allele. In addition to a dTAG fusion (Nabet et al., 2018) for controlled protein degradation, this allele contains an HA tag fusion to enable immunoprecipitation (IP) and immunofluorescence (IF) independent of the availability of TF-specific antibodies, and a transcriptional reporter encoding a nuclear-localized EGFP (**Fig.1C**). Flow cytometric analysis revealed a low level of *Otx2* expression in mESCs, but significant upregulation in derivative EpiLCs and epiblast stem cells (EpiSCs) (**Fig. S1C**), representing two successive stages of the post-implantation epiblast before germ layer specification. This observation is consistent with previous results (Buecker et al., 2014) and *Otx2* expression kinetics *in vivo* (Acampora et al., 2013). Treatment of OTX2-dTAG EpiLCs with the inducer molecule dTAG-13 resulted in reproducible and highly efficient (>90%) degradation of OTX2 within one hour (h) of culture (**Fig.1D**), documenting our ability to acutely control OTX2 levels.

### OTX2 depletion impairs DE formation from mouse pluripotent stem cells

To explore the role of OTX2 during differentiation, we converted EpiLCs derived from OTX2-dTAG mESCs into anterior primitive streak derivatives that emerge during gastrulation using micropattern technology (Morgani et al., 2018)(**Fig. S1D**). Immunostaining revealed a quantitative reduction in the number of cells expressing the DE marker SOX17 (Kanai-Azuma et al., 2002; Morgani et al., 2018) after depletion of OTX2 at the onset of differentiation. In contrast, FOXA2^+^SOX17^-^ putative axial mesoderm cells were still efficiently formed (**Fig.1E,F**). We also observed a slight increase in the levels of the epiblast marker SOX2 with loss of OTX2 (**Fig.1E,F**). These observations demonstrate perturbed gastruloid formation without OTX2 and suggest a previously unappreciated and specific role of this TF for DE development.

To further dissect the molecular function of OTX2 during DE development, we implemented a recently published directed differentiation approach which efficiently converts EpiSCs into primitive streak (PS) (via transient WNT activation) followed by DE (via BMP inhibition and high levels of Activin A/Nodal signaling) (Medina-Cano et al., 2022) (**Fig.1G**). We established and validated stable homozygous OTX2-dTAG EpiSC lines and differentiated them towards DE, confirming expression of canonical protein markers OCT4 (EpiSC/PS), T(PS) and SOX17 (DE) by the majority (>90%) of cells at the expected timepoints (**Fig. S1E**). *Otx2* expression (measured by *Otx2*-EGFP or HA IF) was detected in >95% of cells at all stages and gradually increased in intensity with highest levels attained in DE (**Fig.1H**), but OTX2 levels were reduced to background after 1h of culture of PS or DE in presence of dTAG-13 (**Fig. S1F**). OTX2 depletion at the PS stage (i.e., concurrent with DE specification) (**Fig.1G**) resulted in reduced differentiation into DE (CXCR4^+^PDGFRA^-^ cells), with a concordant increase in PDGFRA^+^ putative early mesodermal cells (Takenaga et al., 2007) (**Fig. S1G** and **Fig.1I**). In addition, we observed reduced levels of the DE-associated TF SOX17 (**Fig.1J**). This effect was specific to OTX2-dTAG cell lines treated with degrader, as dTAG-13 treatment did not affect SOX17 expression in parental (WT) cells (**Fig. S1H**) and DE differentiation was unaffected by DMSO treatment (vehicle control) in OTX2-dTAG cells (**Fig. S1G**).

These findings suggest that OTX2 is important for differentiation towards DE and, possibly, away from mesoderm. OTX2 works in a partially cell non-autonomous manner during anterior neuroectoderm induction (Rhinn et al., 1999). However, analysis of DE established after mixing different ratios of OTX2-dTAG and parental (WT) EpiSCs, revealed no evidence for rescue of CXCR4 levels in OTX2-depleted cells by WT cells or reduced CXCR4 levels in WT cells in presence of OTX2-depleted cells (**Fig. S1I**). These observations support the cell-autonomous functions of OTX2 in driving DE-associated gene expression.

### OTX2 loss affects specific developmental programs during DE specification and maintenance

To characterize transcriptional consequences of OTX2 depletion during DE specification on a genome-wide scale and at single cell resolution, we applied single cell RNA sequencing (scRNA-seq) analysis (10X Genomics platform). For this, we used cells generated with the directed differentiation paradigm described above. Multiplexed samples – EpiSCs, DE derived in presence of DMSO (control DE) and DE after OTX2 depletion in PS when induced to become DE (OTX2^depl_PS^ DE) – were subjected to standard processing and QC procedures including elimination of background reads and low-quality cells (Fleming et al., 2022) and analyzed with Scanpy (Wolf et al., 2018). Both control and OTX2 depleted DE clustered away from EpiSCs and showed extinction of pluripotency-associated transcripts, absence of markers for visceral (*Sox7*) and primitive (*Ttr*) endoderm, as well as high levels of a subset of DE-specific markers such as *CD24a*, *Larp7* and *Hhex* (Moore et al., 2014; Pijuan-Sala et al., 2019; Wang et al., 2012) (**Fig. S2A**). We also did not observe differences in cell cycle state based on expression of the G2/M indicators *Aurka* and *Plk1* (Liu et al., 2022) (**Fig. S2A**).

Nevertheless, OTX2^depl_PS^ DE formed a distinct cluster on the UMAP projection (**Fig.2A** and **Fig. S2B**), suggesting that OTX2 depletion redirects the developmental trajectory of DE formation from the epiblast and results in a distinct transcriptional state. Accordingly, we identified a total of 1,646 differentially expressed genes (DEGs) (RNA score>1;logFC>1;padj<0.05) between control and OTX2^depl_PS^ DE (**Table S2**), similar fractions of which were downregulated (44.9%) or upregulated (55.1%) (**Fig.2B**). Gene ontology analysis of DEGs suggested dysregulation of WNT signaling, cell adhesion/migration and cell differentiation in absence of OTX2 (**Fig.2C**). At the gene level, we observed downregulation of DE-associated TFs (such as *Sox17* and *Hesx1*), antagonists of WNT signaling (*Dkk1*, *Shisa2*) and cell migration/adhesion regulators (*Cdh1*, *Sema6d* and *Emb*) in OTX2^depl_PS^ DE (**Fig.2D** and **Fig. S2C**). Upregulated genes included agonist and canonical targets of WNT and FGF/regulators (*Cdh2*, *Sema3a*) (**Fig.2D** and **Fig. S2C**).

**Figure 2.**
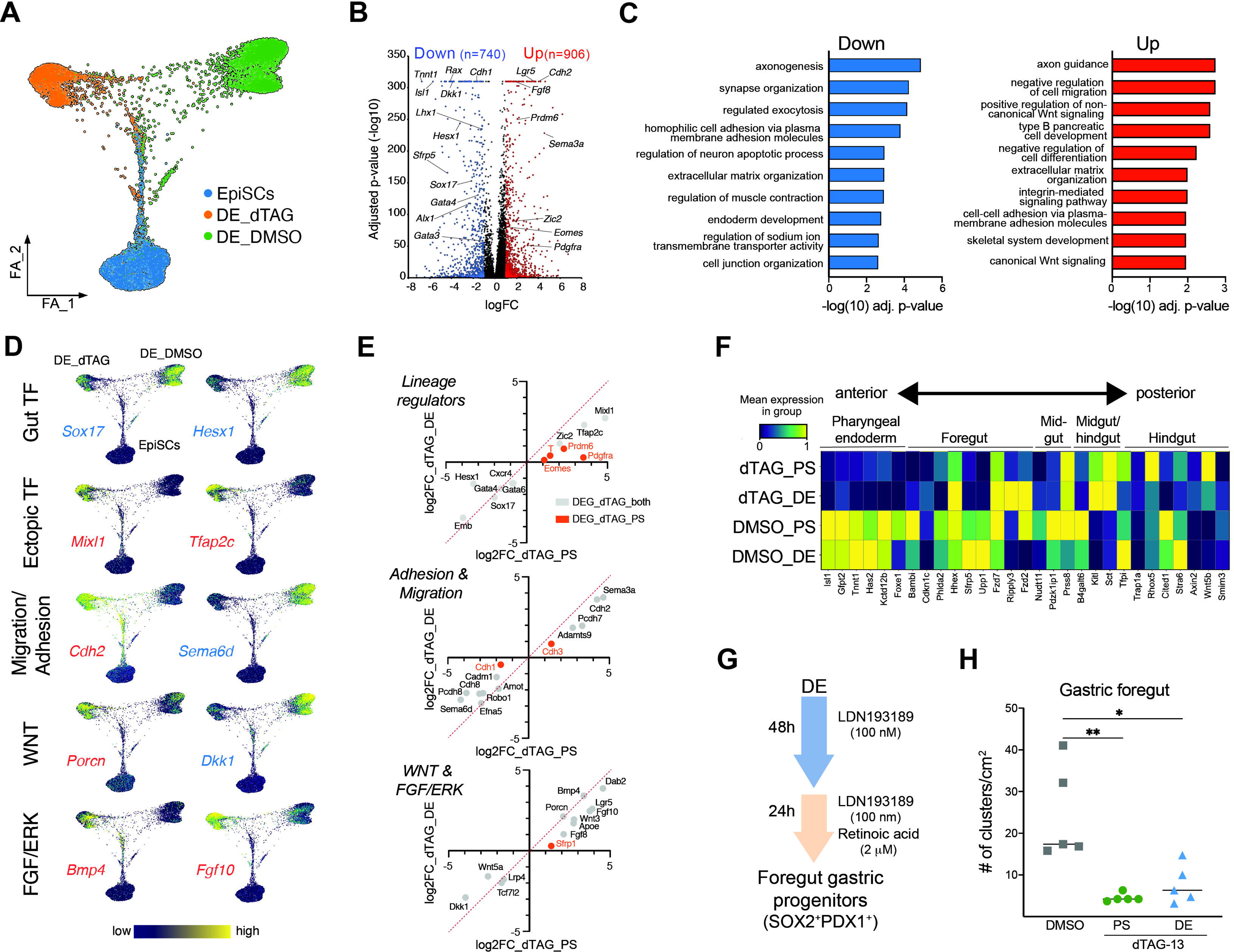
Transcriptional dysregulation in DE upon OTX2 depletion. A. Force-directed layout graph of scRNA-seq analyses on WT EpiSCs, DE treated with DMSO and DE treated with dTAG-13 at PS (OTX2^depl_PS^) DE cells. B. Volcano plot showing expression changes (logFC) and statistical significance (padj) in OTX2^depl_PS^ DE with DEGs (logFC>1;padj<0.05) highlighted in blue (DEG^DOWN^) and red (DEG^UP^), respectively. DEGs highlighted in the text or in other figure panels are annotated. C. Gene ontology (GO) analysis for DEG^DOWN^ and DEG^UP^ in OTX2^depl_PS^ DE using ENRICHR, listing top ranked GO terms for each gene category. D. Normalized expression profiles of select DEG^DOWN^ (blue names) and DEG^UP^ (red names) in OTX2^depl_PS^ DE that associated with indicated cellular programs. E. Comparison of the fold-change effect (dTAG-13:DMSO) on expression of select DEGs when OTX2 is depleted during (dTAG_PS; x-axis) or after (dTAG_DE; y-axis) DE specification. Genes DEGs only upon OTX2 depletion at PS are highlighted in red, all other genes are DEGs in both conditions. F. Normalized expression levels of a panel of anterior-posterior gut tube markers (Pijuan-Sala et al., 2019) OTX2^depl_PS^ DE, OTX2^depl_DE^ DE and in control DE treated with DMSO. G. Protocol to differentiate DE into SOX2^+^PDX1^+^ gastric foregut progenitors. H. Quantification of SOX2^+^PDX1^+^ foregut clusters formed after OTX2 depletion at either PS or DE compared to cells differentiated in DMSO. (*)p<0.05 and (**)p<0.01 with one-way ANOVA and Kruskal-Wallis multiple comparison test.

Of note, we also observed significant upregulation of several TFs (*Prdm6*, *Mixl1* and *Tfap2c*) (**Fig.2D** and **Fig. S2C**) associated with other embryonic lineages emerging in early post-gastrulation embryos, including mesoderm, neuroectoderm and primordial germ cells (PGCs) (**Fig. S2D**). We also confirmed upregulation of mesoderm associated *Pdgfra* (**Fig. S2E**). Importantly, ectopic lineage markers were co-expressed in OTX2^depl_PS^ DE at the single cell level (**Fig. S2E**), suggesting that OTX2 loss does not result in the emergence of multiple distinct lineages but rather to partial derepression of specific non-endodermal markers in the same cells. Together, these observations demonstrate that loss of OTX2 during DE specification results in the dysregulation of specific gene expression programs and a profound redirection of DE identity. We confirmed dysregulation of major genes representing the affected programs (TFs, WNT and FGF/signaling and cell adhesion/migration) upon OTX2 depletion at PS in an independent cell line (**Fig. S2F**).

To compare molecular consequences when losing OTX2 before or after endodermal identity has been established (Medina-Cano et al., 2022), we depleted OTX2 24h after initiation of DE specification, followed by scRNA-seq analysis. This strategy resulted in a similar number of DEGs (n=1,477) than depletion at PS (**Fig. S2G**) with overall concordant gene expression changes triggered by OTX2 loss before and after DE specification (**Fig.2E**). For example, hallmark molecular changes observed upon OTX2 loss at PS – such as dysregulated expression of WNT/FGF signaling components and of cell adhesion/migration regulators as well as reduced levels of DE markers – were also observed in OTX2^depl_DE^ DE (**Fig.2E**). However, elevated expression of genes associated with nascent mesodermal lineages such as *Eomes*, *T*, *Prdm6* and *Pdgfra* was more pronounced in OTX2^depl_PS^ DE (**Fig.2E**), suggesting that a repressive role of OTX2 at these loci may no longer be required once DE has been specified.

Many of the shared gene expression changes induced by OTX2 depletion before or after DE specification – such as the observed evidence for increased WNT and FGF/ERK signaling (Loh et al., 2014; Pan et al., 2020) – are consistent with loss of anterior-posterior (AP) identity. Indeed, evaluation of the relative expression levels of a panel of marker genes for different endodermal lineages along the AP axis (Pijuan-Sala et al., 2019) revealed a pronounced deficit in the expression of specific anterior markers in the absence of OTX2 (**Fig.2F**). Accordingly, further differentiation of OTX2-depleted DE towards the foregut lineage (**Fig.2G**) revealed an impairment to form clusters of SOX2^+^PDX1^+^ gastric progenitors (Medina-Cano et al., 2022). Evidence for altered foregut development were observed independently of whether OTX2 was depleted at the PS stage or after DE specification (**Fig.2H** and **Fig. S2H**). These molecular and functional analyses document that OTX2 contributes to establishing and maintaining gene expression programs required for successful anteriorization of the developing DE.

### OTX2 drives locus-specific chromatin remodeling during DE specification

Having established a requirement of OTX2 for proper specification and anterior patterning of DE, we next sought to investigate the mechanisms of gene regulation by OTX2 during DE specification. To identify potentially direct OTX2 targets in EpiSCs and DE, we conducted CUT&RUN experiments with a validated antibody (Alexander et al., 2018; Seah et al., 2019) against the HA epitope that is part of our multipurpose degron allele (**Fig.1C**). CUT&RUN revealed 1,286 high-confidence OTX2 binding sites (see Methods section) that were unique to undifferentiated EpiSCs (“EPI peaks”), 5,035 binding sites shared between EpiSCs and DE (referred to as “primed DE peaks” hereafter) and 37,676 OTX2 bindings unique to DE 24h after induction from PS (referred to “*de novo* DE peaks”) (**Fig.3A**). In contrast, HA CUT&RUN in wildtype DE (i.e., not carrying the HA epitope) revealed only background signal (**Fig. S3A**), supporting the specificity of OTX2 detection. The drastic increase in the number of OTX2 binding sites – and the elevated strength of primed peaks from EpiSCs to DE (**Fig. S3B**) – coincides with the marked transcriptional upregulation of *Otx2* during DE specification (see **Fig.1H**) and indicates extensive reorganization of the OTX2 cistrome during this developmental transition.

**Figure 3.**
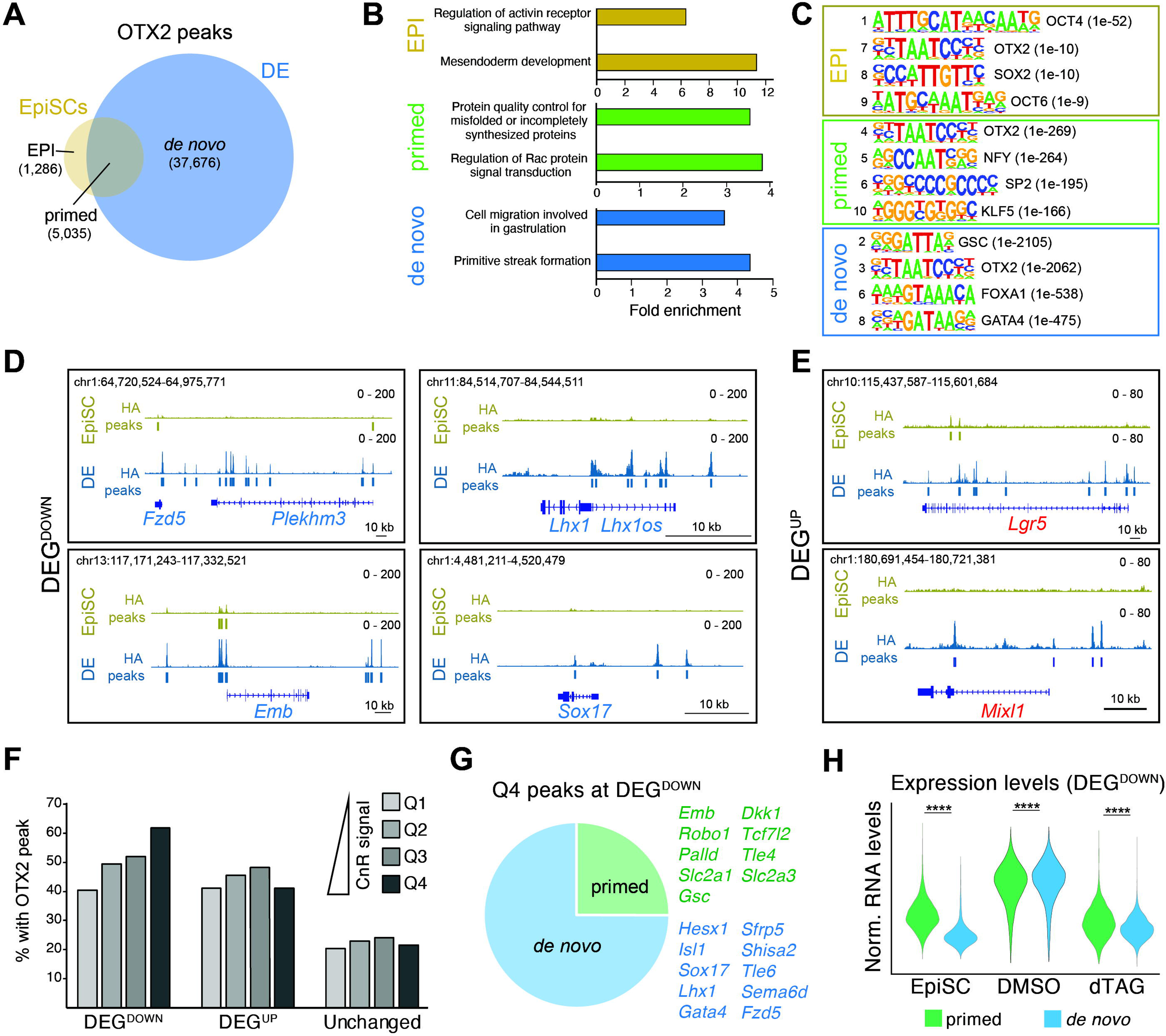
Dynamics of OTX2 genome occupancy during DE specification. A. Classification and number of OTX2 binding sites identified by CUT&RUN in EpiSCs and DE. B. GO analysis of genes in proximity of EPI, primed, and *de novo* OTX2 binding sites. C. HOMER TF motif analysis at EPI, primed, and *de novo* OTX2 binding sites. The analysis was restricted to TFs expressed in DE and/or EpiSCs. D. IGV tracks of OTX2 genome occupancy at select DEG^DOWN^ loci with examples of genes exhibiting primed (left two panels) or *de novo* (right two panels) OTX2 binding. E. IGV tracks of OTX2 genome occupancy at select DEG^UP^ loci with examples of primed (top panel) and *de novo* (bottom panel) OTX2 binding. F. Percentage of DEG^DOWN^, DEG^UP^ and gene loci unaffected by OTX2 depletion (“unchanged”) with OTX2 peaks of increasing (Q1 to Q4) intensity. G. Distribution of DEG^DOWN^ gene loci with evidence of primed (site bound both in EpiSCs and DE) and *de novo* (site bound only in DE) OTX2 binding with example genes highlighted. Note that all gene loci with primed peaks also harbor additional *de novo* peaks but not vice versa. H. Normalized expression levels of DEG^DOWN^ with either primed or *de novo* only OTX2 binding in EpiSCs, DE_DMSO an DE_dTAG. (****)p<0.0001 with two-sided unpaired Wilcoxon test.

EPI and *de novo* DE peaks both were most frequent at intronic and distal intergenic sites (**Fig. S3C**) and enriched for genes associated with stage-specific developmental functions, such as Activin receptor signaling and multilineage differentiation (EPI peaks) or cell migration and primitive streak formation (DE peaks) (**Fig.3B**). For example, EpiSC-specific binding was evident at *Pou5f1*, a pluripotency associated locus previously reported to be regulated by OTX2 (Di Giovannantonio et al., 2021) (**Fig. S3D**), and occurred at sites enriched for binding motifs of pluripotency-associated TFs (OCT4, SOX2, OCT6) (**Fig.3C**). In contrast, *de novo* peaks were enriched for binding motifs of TFs with known functions in endoderm development (such as GSC, FOXA1, GATA4) (**Fig.3C**). Primed DE peaks were frequently promoter proximal (**Fig. S3C**) and enriched for genes associated with universal cellular processes such as protein quality control and Rac signaling (**Fig.3B**) as well as for motifs of non-cell type specific TFs (**Fig.3C**), possibly reflecting their position within promoters. These observations document that OTX2 occupies candidate regulatory elements of critical lineage-associated genes in a stage-specific manner, likely in collaboration with other developmental TFs.

We observed well-defined OTX2 peaks at many DEG^DOWN^ and DEG^UP^, representing the major cellular programs transcriptionally dysregulated upon OTX2 depletion (**Figs.3D,E** and **Table S3**). To systematically investigate the relationship between OTX2 binding and the transcriptional responses triggered by its loss, we analyzed frequency, strength, genomic positioning, and developmental dynamics of OTX2 peaks at genes affected by OTX2 depletion. To reduce the impact of lowly expressed genes on this analysis, we focused on DEGs with an RNA score>10, covering 448 DEG^UP^ and 393 DEG^DOWN^ (**Table S2**). As reference, we used 1,169 gene loci expressed in DE but unaffected (padj>0.9) by OTX2 depletion. In accordance with a direct role of OTX2 in controlling DE transcription) we observed that >80% of DEG^DOWN^ and >75% of DEG^UP^ but only slightly less than 50% of unaffected loci were bound at least once by OTX2 at their promoters or distal regions(**Fig. S3E**). OTX2-bound DEGs on average also showed a significantly higher number of OTX2 peaks than unaffected genes (**Fig. S3F**). In a clear distinction, OTX2 peaks at DEG^DOWN^ were significantly stronger than peaks at DEG^UP^ and at unaffected genes (**Fig. S3G** and **Fig.3D,E**). We also observed a pronounced enrichment of DEG^DOWN^ but not of DEG^UP^ in vicinity of the strongest peaks (4^th^ quartile; Q4) (**Fig.3F**). This suggests that the locus-specific regulatory function of OTX2 (activator versus repressor) may in part be determined by the affinity of its target cis regulatory elements, as has been proposed for homeobox TFs in other developmental contexts (White et al., 2016).

Despite the overall low abundance of primed peaks (11.8%) among all DE-associated peaks (**Fig.3A**), more than a quarter (26.3%) of DEG^DOWN^ showed evidence for OTX2 binding in EpiSCs (**Fig.3G**) with almost all DE-associated loci that were already bound by OTX2 in EpiSCs acquiring additional peaks upon differentiation (**Fig. S3H** and **Fig.3D,E**). Consistent with a priming function of OTX2 before DE differentiation, genes with primed peaks were already expressed at higher levels in EpiSCs before becoming further upregulated during DE specification (**Fig.3H**). Both primed and *de novo* bound loci enriched for genes encoding regulators of cell adhesion and migration (such as *Emb*, *Robo1* and *Sema6d*) and antagonists of WNT signaling (*Dkk1* and *Shisa2*) (**Fig.3G**). In contrast, DEG^DOWN^ loci encoding TFs regulating DE and derivative anterior lineages such as *Hesx1*, *Sox17* and *Isl1* were bound by OTX2 in DE only (**Fig.3D,G** and **Table S3**). These observations are consistent with the notion that OTX2 binding in the epiblast primes transcriptional programs broadly required for gastrulation and germ layer formation. The full activation of OTX2-controlled DE-specific genes, however, requires additional regulatory remodeling upon receipt of developmental signals.

### OTX2 functions partially by controlling chromatin accessibility at development loci

We next examined to what degree stage specific OTX2 binding impacts chromatin accessibility during DE differentiation. When focusing on differentially accessible regions (DARs) between EpiSCs and DE (Medina-Cano et al., 2022), we noticed that more than 40% of DE-specific DARs (n=20,475) overlapped with OTX2 target sites (either primed or *de novo*) (**Fig. S4A**), while the vast majority (∼90%) of EpiSC DARs did not. To test experimentally whether OTX2 binding actively contributes to chromatin opening during endoderm specification, we performed ATAC-seq analysis of DE 24h after OTX2 depletion at the PS stage. This revealed 3,381 high-confidence DARs (logFC>1;padj<0.05) compared to DMSO-treated controls. Among DARs, 26.1% showed elevated and 73.9% reduced ATAC-seq signal (**Fig.4A** and **Fig. S4B**), demonstrating that OTX2 favors the establishment or maintenance of accessible chromatin in DE. The majority but not all chromatin changes induced by OTX2 depletion occurred at sites normally undergoing accessibility changes during DE specification (**Fig. S4C**).

**Figure 4.**
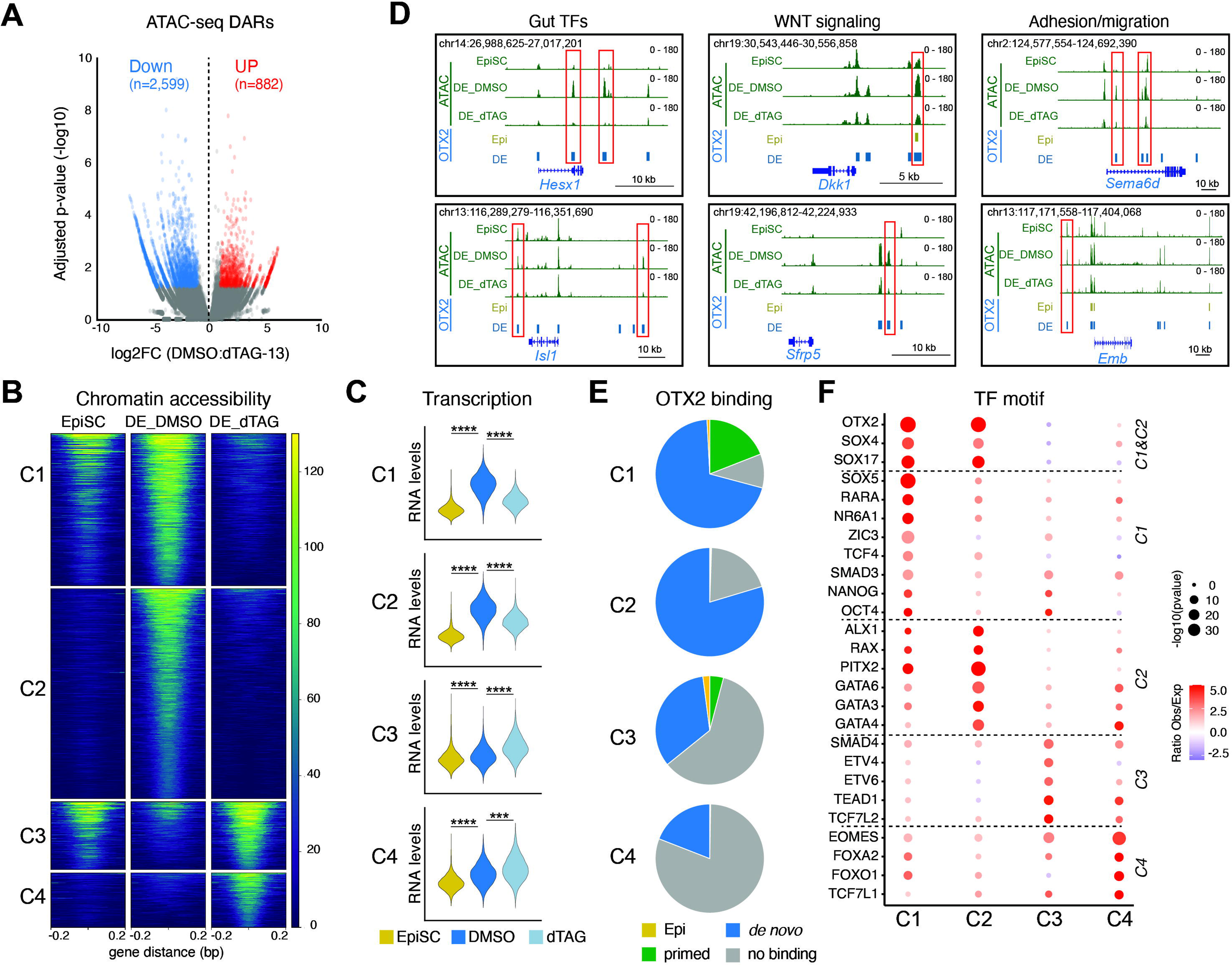
OTX2-dependent remodeling of chromatin accessibility during DE specification. A. Volcano plot showing ATAC-seq signal in DMSO-treated DE versus DE treated with dTAG-13 at PS. Significantly different DARs (logFC>1;padj<0.05) are colored in blue (DAR^DOWN^) and red (DAR^UP^). B. Four categories of DARs (C1-C4) identified by supervised clustering of ATAC-seq signal intensity in EpiSCs, control DE and OTX2-depleted DE. C. Normalized expression levels of genes in vicinity of C1 to C4 DARs in indicated samples. (***)p<0.001 and (****)p<0.0001 with two-sided unpaired Wilcoxon test. D. Example of chromatin accessibility changes at DEG^DOWN^ loci representing major cellular programs affected by OTX2. E. Frequency and type of OTX2 binding at C1 to C4 DARs. F. HOMER TF motif enrichment at C1-C4 DARs. Analysis was restricted to TFs expressed in EpiSCs, DE, or both. Dotted lines indicate TFs exhibiting enrichment at similar DAR categories.

Supervised clustering of all lost or gained DARs upon dTAG-13 treatment in DE along with their accessibility levels in wildtype EpiSC, generated four distinct ATAC-seq peak clusters (C1-C4). C1 genomic sites (n=1,091) were accessible in EpiSCs and DE but lost accessibility in DE upon OTX2 depletion, whereas C2 sites (n=1,508) normally gained accessibility in DE but failed to do so in absence of OTX2 (**Fig.4B**). Both C1 and C2 DARs were enriched at promoter distal regions (**Fig. S4D**) and showed strong, OTX2-dependent transcriptional upregulation of associated genes during DE specification (**Fig.4C**). In total, 32.6% of DEG^DOWN^ were associated with local loss of chromatin accessibility upon OTX2 depletion (**Fig. S4E**), including gene loci encoding negative regulators of WNT signaling (*Dkk1, Sfrp5*), cell type-specific transcription (*Hesx1, Isl1*) and cellular migration/adhesion (*Emb, Sema6d*) (**Fig.4D** and **Table S4**). Importantly, most C1 and C2 DARs were bound by OTX2 (**Fig.4E**) with strongest binding at C1 (**Fig. S4F**). These observations support a direct role of OTX2 in controlling chromatin accessibility and transcriptional output at these loci. The regulatory function of OTX2 at a subset of C1 sites includes preoccupancy in EpiSCs (**Fig.4E**). Of note, while binding motifs of OTX2 and SOX factors were enriched at both C1 and C2 DARs (**Fig.4F and S4G**), TCF4, NR6A1 and pluripotency factors NANOG and OCT4 were predominantly associated with C1 DARs only and GATA factors with C2 DARs only (**Fig.4F and S4G**). These observations suggest that OTX2-dependent chromatin opening during DE specification enables binding of additional, stage-specific co-regulators.

As mentioned above, OTX2 depletion also resulted in gain of chromatin accessibility (C3 and C4 DARs). C3 sites (n=489) were pre-accessible in EpiSCs and failed to appropriately close in DE upon OTX2 depletion, whereas C4 sites (n=393) were normally closed in both EpiSCs and DE but aberrantly gained accessibility in DE in absence of OTX2 (**Fig.4B**). Concordantly, genes associated with C3 and C4 peaks showed transcriptional upregulation upon OTX2 depletion (**Fig.4C**). Strikingly, the majority of C3 and C4 DARs did not overlap with OTX2 binding sites (**Fig.4E**), suggesting indirect activation of most of these loci downstream of OTX2 loss. Moreover, even C3/C4 DARs bound by OTX2 were not enriched for the OTX2 consensus motifs (**Fig. S4G**), supporting the notion that binding of OTX2 to loci it represses is mechanistically distinct from OTX2 binding to loci this TF activates. C3/C4 DARs were enriched for a distinct set of regulators than C1/C2 DARs, including the endomesodermal TFs EOMES and FOXA2 and the WNT mediators TCF7L1/2 (**Fig.4F and S4G**). This observation suggests that OTX2 loss increases levels and/or activity of these factors, resulting in ectopic chromatin activation. Together, these observations show that OTX2 engages in both activating and repressive functions during DE specification and exerts part of its activator function by maintaining pre-existing and establishing new chromatin accessibility at distal gene regulatory elements that control the expression of endoderm-associated genes.

### OTX2 is required for faithful specification of human DE

Compared to most other TFs associated with endodermal differentiation, OTX2 has an unusual high degree (99.7%) of amino acid sequence conservation between mouse and human (Cunningham et al., 2022) (**Fig. S5A**). This suggests evolutionary pressure to preserve protein function and establishes a unique opportunity to study shared and species-specific cis-regulatory aspects of transcriptional control during endoderm specification. To determine the role of OTX2 during human DE development, we established homozygous human embryonic stem cell (hESC) lines carrying the identical multipurpose degron cassette we employed in mouse (**Fig.5A**). For this, we used MEL-1 and H9 hESCs, two well-characterized lines with established DE potential (Chia et al., 2019; Jiang et al., 2013). In both parental backgrounds, the hOTX2-dTAG allele allowed rapid (1h) and efficient (>90%) OTX2 depletion in hESC and derivative DE (**Fig.5B**), using the degrader dTAG-v1 (Nabet et al., 2020) which we found more effective in human cells than dTAG-13 (**Fig. S5B**).

**Figure 5.**
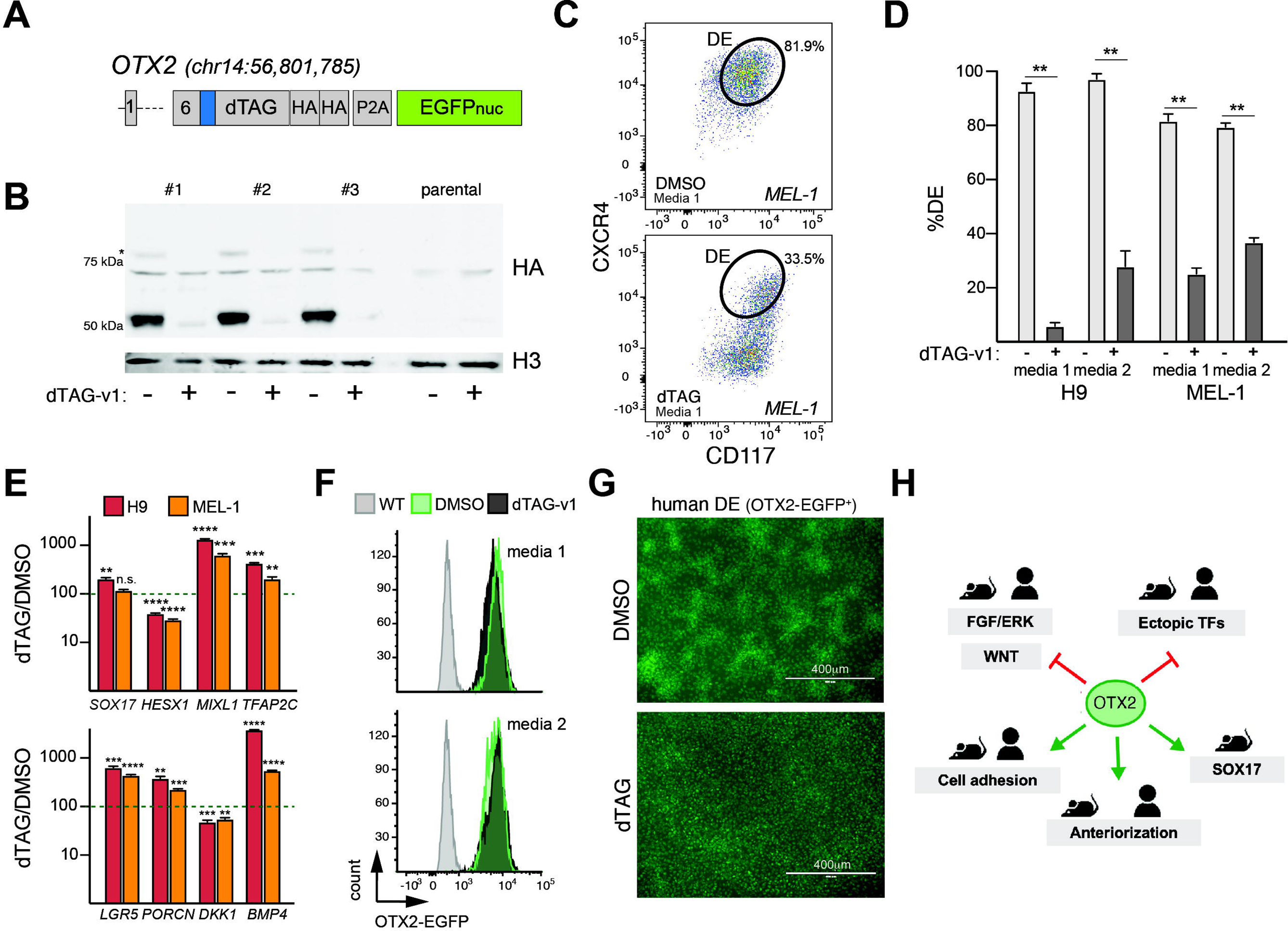
Impaired human DE formation upon OTX2 depletion. A. Design of the human OTX2-dTAG allele. Coordinates indicate the position of the *Otx2* STOP codon (in hg38) replaced with the multipurpose cassette. The blue box indicates a linker peptide. B. Anti-HA WB with OTX2-dTAG hESCs (n=3 lines) and parental cells after 1h of culture in the presence of dTAG-v1 or DMSO. * = minor band indicating incomplete cleavage of P2A fusion protein. C. Representative flow cytometry plots showing CXCR4 and CD117 expression in human DE derived from MEL-1 cells after culture in the presence of DMSO (top) or dTAG-v1 (bottom) from the PS stage onwards. Cells with a canonical DE cell surface phenotype are highlighted. D. Abundance of cells with the canonical DE phenotype (CXCR4^+^CD117^+^) in cultures initiated with two independent OTX2-dTAG hESC lines and exposed to either DMSO or dTAG-v1 in two different culture conditions (n=3 independent cultures). (***)p<0.001 or (****)p<0.0001 with multiple T-tests and Bonferroni-Dunn correction. E. Measurement by qPCR of effect of dTAG-v1 treatment on select human genes whose mouse homologues are altered in their expression levels by OTX2 depletion in DE. (*)p<0.05, (***)p<0.001 or (****)p<0.0001 with multiple T-tests and Bonferroni-Dunn correction. F. EGFP fluorescence levels in DE cultures established in either presence of DMSO or dTAG-v1. Cultures established from parental non-transgenic (WT) human ESCs serve as control for background fluorescence. G. Representative low-magnification fluorescent live cell images of OTX2-EGFP^+^ human DE derived in presence of DMSO (top) or dTAG-v1 (bottom). Images are of MEL-1 derived cells using Protocol 1, but similar differences in cell clustering were observed with H9-derived cells and with DE generated using media 2. H. Schematic highlighting major functions of OTX2 during mouse and human DE specification.

We differentiated OTX2-dTAG hESCs towards DE with two slightly different protocols – one using identical signaling manipulations to the one applied in mouse (“Protocol 1”) (Medina-Cano et al., 2022), the other applying additional PI3K inhibition during PS specification (“Protocol 2”) (Loh et al., 2014) (**Fig. S5C**). In accordance with the developmental resemblance of human pluripotent cells to the post-implantation epiblast, we observed widespread *OTX2* expression in OTX2-dTAG hESCs (**Fig. S5D**), mirroring the situation in mouse EpiSCs and EpiLCs (**Fig. S1C**). *OTX2* expression levels strongly increased during commitment to human DE (**Fig. S5E**), as we also had observed in mouse (**Fig.1H**), suggesting similarities in the manner the *OTX2* locus is controlled by external signaling cues in both species.

Differentiation of OTX2-dTAG hESCs towards endoderm in the absence of dTAG-v1 yielded predominantly cells co-expressing the DE markers CXCR4 and CD117 (**Fig.5C,D**), demonstrating that the degron allele does not interfere with endodermal differentiation. In contrast, depletion of OTX2 from the PS stage onwards resulted in significantly fewer CXCR4^+^CD117^+^ cells (**Fig.5C,D**). Analysis by qPCR of select gene loci representing pathways and processes that are affected by OTX2 depletion in mouse DE, revealed evidence for dysregulation of similar developmental modules. Thus, we observed altered expression of developmental TFs, including lower levels of the DE-associated TF *HESX1* and elevated levels of “ectopic” TFs such as *MIXL1* and *TFAP2C* in human DE specified in the absence of OTX2 (**Fig.5E**). In contrast, SOX17 levels were not downregulated at RNA or protein level in human DE upon OTX2 depletion (**Fig.5E** and **Fig. S5F**), suggesting differences in the regulation of this locus between mouse and human DE (see **Fig.1J** and **Fig.2D**).

Expression changes of genes encoding WNT and FGF/ERK signaling components changed in a manner consistent with activation of these two pathways in human DE in the absence of OTX2 (**Fig.5E**), similar to what we had observed in mouse DE. While levels of the transcriptional OTX2-EGFP reporter were only moderately affected by OTX2 depletion (**Fig.5F**), fluorescence microscopy showed that EGFP^+^ human DE lacking OTX2 protein failed to form the dense cell clusters that were abundant in endodermal cultures expressing this TF (**Fig.5G**). This suggests a deficit in the ability of OTX2-deficient DE to self-organize that is reminiscent of the observations made in mouse foregut (see **Fig.2H and S2H**). Together, these observations are consistent with the notion that OTX2 is an important regulator of human gut tube specification and suggest that specific developmental functions of this TF are conserved between mice and humans (**Fig.5H**).

## Discussion

To facilitate studying TF function during mammalian germ layer specification, we implemented an experimental platform that combines efficient and scalable *in vitro* models of embryonic lineage specification with genetic tools for controlled protein degradation. Taking this generalizable approach, we have identified and characterized a previously unappreciated role of OTX2 in DE specification and patterning. In contrast to other TFs regulating endoderm development such as SOX17 (Kanai-Azuma et al., 2002) or the early endomesodermal specification factor EOMES (Arnold et al., 2008) whose loss in KO mouse models results in entire parts of the early gut tube missing, OTX2 depletion in our system does not result in a complete loss of endoderm identity or the generation of non-viable cells. Rather, acute degradation of OTX2 during or after DE formation leads to incorrect patterning with OTX2-depleted DE characterized by dysregulation of WNT and FGF/ERK signaling, altered expression of migratory and cell adhesion molecules, as well as ectopic expression of regulators of other early embryonic lineages. These abnormalities prevent the faithful establishment of anterior endoderm identity. Our results are in accordance with the mislocalization of endomesodermal cells and impaired foregut formation in OTX2 KO mice (Acampora et al., 1995; Kimura et al., 2000).

At the molecular level, OTX2 binds to distal gene regulatory elements at loci encoding regulators of cellular signaling, cell adhesion, and gut tube development such as *Dkk1*, *Cdh1*, *Hesx1*, and *Isl1*. At these loci, OTX2 is required to establish and maintain chromatin accessibility. These observations establish OTX2 as a direct regulator of – and potential pioneer factor for – a subset of candidate enhancers involved in DE specification and patterning. Of note, at several genes encoding known regulators of gastrulation and/or gut tube development, OTX2 binding precedes transcriptional upregulation. This suggests a priming function of OTX2 binding in the epiblast, reminiscent of the pre-positioning reported for other TFs, such as OCT2, during lymphocyte development (Doane et al., 2021). The changes in OTX2 genome binding we observed between epiblast and endoderm continue the genome occupancy dynamics initiated when this TF becomes activated upon exit from naïve pluripotency (Navarra et al., 2016; Yang et al., 2014). While requiring further investigation, it is plausible that *de novo* OTX2 binding in DE is driven partly by the availability of different co-factors (resulting in a shift away from sites occupied by pluripotency-associated factors such as OCT4 and towards sites occupied by endoderm regulators such as SOX2/SOX17 and GATA factors) and partly by the elevated OTX2 levels in DE (possibly resulting in occupation of low-affinity binding sites). The observation that only comparatively few OTX2-bound genes are differentially expressed upon OTX2 loss might suggest promiscuous binding or redundancy with another TF. On the other hand, not all OTX2-regulated genes (i.e., bound by OTX2 and downregulated in dTAG conditions) lose chromatin accessibility upon OTX2 depletion. This points to additional mechanisms beyond chromatin opening that OTX2 may employ to drive gene expression, such as changes in enhancer activity and/or chromatin looping (Uyehara and Apostolou, 2023).

We observed aberrant chromatin opening upon OTX2 depletion at only a subset of gene loci upregulated in the absence of OTX2, suggesting that gene repression by this TF is primarily mediated by a mechanism not affecting chromatin accessibility. Of note, studies in frogs have suggested that OTX2 can directly exert both activating and repressive functions, depending on the nature of cis-regulatory elements it binds to and the co-factors it engages with (Yasuoka et al., 2014). In agreement with this, we observed weaker binding of OTX2 to gene loci it represses and no enrichment of the OTX2 consensus motif at sites gaining chromatin accessibility in the absence of this TF. We surmise that OTX2 achieves gene repression in DE by a variety of direct and indirect mechanisms, including counteracting the expression and/or activity of TFs driving alternative lineages and suppression of WNT signaling.

Our stage-specific deletion experiments demonstrate a requirement for OTX2 beyond DE specification. In fact, most genes and developmental programs affected in DE upon OTX2 loss before or after endoderm specification are shared. Intriguingly, high levels of *Otx2* expression are maintained at least until the anterior foregut and possibly beyond in thyroid and thymus progenitors (Nowotschin et al., 2019), raising the possibility that OTX2 functions throughout the progressive anteriorization of the developing gut tube. The similar expression kinetics of *Otx2*/*OTX2* during the early stages of mouse and human *in vitro* development, the enrichment of the OTX2 motif at mesendodermal enhancers during hESC differentiation (Tang et al., 2022), and our limited analysis of gene expression changes at murine OTX2 target genes in OTX2-depleted human DE, suggests at least partial conservation of OTX2 function during mammalian evolution. Furthermore, the observation that some of the OTX2 targets we identified in DE are shared with other lineages, including *Dkk1* in the anterior visceral endoderm (Kimura-Yoshida et al., 2005) and *Hesx1* in the forebrain (Spieler et al., 2004), supports the existence of a shared molecular blueprint of AP patterning that employs OTX2. The full repertoire of stage-specific versus tissue-specific versus species-specific versus “universal” OTX2 target genes – and the mechanisms of their regulation by OTX2 – remains to be determined.

Upon OTX2 depletion, we observed elevated expression of some signaling and transcriptional regulators associated with the primitive streak and/or the early mesoderm (*Pdgfra, Fgf8, Mixl1, Eomes, Prdm6*), the germline (*Tfap2c)* and, to a more limited degree, the neuroectoderm (*Zic2, Zic5*). The de-repression of *Tfap2c/TFAP2C*, which encodes a TF required for germline development (Kojima et al., 2021), is consistent with a germline-repressive function of OTX2 in mice (Zhang et al., 2018) and humans (Tang et al., 2022). Based on these observations, we speculate that OTX2 facilitates the timely establishment of an endodermal identity in the anterior primitive streak by selectively targeting master regulators of alternative embryonic cell types rather than broadly suppressing the gene expression programs of these lineages. Of note, in an *in vitro* system of mouse PGC formation, OTX2 depletion favors germline over mesoderm differentiation (Di Giovannantonio et al., 2021). This suggests stage-specific functions of OTX2 in somatic lineage specification beyond neuroectoderm and DE, a notion consistent with the expression pattern of *Otx2* in gastrulation-stage embryos (**Fig.1B**).

An ability of OTX2 to directly counteract posterior development has been suggested by microinjection experiments in the frog (Pannese et al., 1995). While further experiments are required to address such an OTX2 function during mammalian development, our scRNA-seq analysis at the DE stage shows no evidence for the upregulation of hindgut regulators or signature genes. This might suggest that OTX2 primarily acts to activate anterior DE loci or reflects the fact that culture conditions employed here are not permissive for posterior gut tube differentiation. The general notion of OTX2 being a gastrulation and germ layer regulator rather than a pluripotency regulator is consistent with the survival of OTX2 KO mice until early organogenesis (Ang et al., 1996) and the mild and delayed defects in mouse pluripotent stem cell lines upon OTX2 deletion (Acampora et al., 2013; Kinoshita et al., 2020).

In summary, our studies provide insight into the transcriptional regulation of early gut tube patterning by OTX2 and suggest similarities in the molecular control of anteriorization across tissues and species. The integration of multipurpose degron alleles into efficient directed differentiation regimens of pluripotent cells represents a tractable and readily generalizable platform to dissect these similarities further and to study other developmental processes in mammals.

## Acknowledgements

We would like to thank Chaitanya Parikh and John Bugay for help with DNA preparations and genotyping, Rachel Glenn, and Emily Corrigan for advice on mouse DE differentiation, Kaushiki Chatterjee and Alejandra Laguillo-Diego for help with Western Blot analysis, and Miriam Gordillo for advice on human DE differentiation. We are grateful to all members of the Stadtfeld and Apostolou labs for their input on this project and feedback on this manuscript. L.E. was a New York Stem Cell Foundation-Druckenmiller Fellow. This research was supported by the New York Stem Cell Foundation. M.S. was supported by grants from the NIH (R01GM145864, R03NS135564), the Simons Foundation, the Tri-Institutional Stem Cell Initiative (Tri-SCI), and the Bohmfalk Charitable Trust. EA was funded by NIH (5R01GM138635 and 5RM1GM139738).

## Author contributions

L.E. generated and characterized murine cell lines, conducted, and analyzed directed differentiation experiments, prepared cells for genomics assays and assisted in micropattern differentiation. D.M. conducted bioinformatics analyses and assisted with foregut differentiation experiments. C.U. conducted CUT&RUN experiments and assisted in bioinformatics analyses. C.S. conducted micropattern experiments. E.S. conducted ATAC-seq experiments. A.P. assisted with bioinformatic analyses. S.M. assisted in characterizing mouse and human degron cell lines. M.S. and E.G. generated human, validated, and differentiated human degron cell lines. The manuscript was written by M.S. and edited by L.S., C.U., D.M., with input from all authors. M.S. acquired funding. T.E., A.K.H., E.A. and T.V. advised on experimental design, provided reagents, and supervised experiments.

## Tables with titles and legends

Table S1. Expression levels of TFs in EpiLCs and mESCs Table S2. DEGs identified by scRNA-seq

Table S3. List of OTX2 peaks identified by HA CUT&RUN Table S4. List of DARs identified by ATAC-seq

Table S5. List of DNA oligos used in this study

## Resource availability Lead Contact

Requests for resources and reagents should be directed to and will be fulfilled by the lead contact, Matthias Stadtfeld.

## Materials availability

Cell lines generated in this study are available upon request from the lead contact.

## Data and code availability

CUT&RUN, scRNA-seq and ATAC-seq data have been deposited at Gene Expression Omnibus (GEO) with accession codes: GSE254428, GSE254431 and GSE254590, respectively. The deposited data will be publicly available as of the publication date.

## Experimental model and subject details

### Mouse cell lines

Parental mouse ESC lines used for gene targeting were KH2 (Beard et al., 2006) or 5.8 (Zhong et al., 2023), both on a C57BL/6J x 129S1 F1 background. OTX2-dTAG mESCs were generated using CRISPR/Cas9. Homology arms covering 778bp upstream and 680bp downstream of the *Otx2* C-terminus were PCR-amplified from genomic DNA and cloned with the FKBP12^F36V^-2xHA-P2A-NLS-EGFP construct into pBluescript (Stratagene) using Gibson assembly. The OTX2-FKBP12^F36V^-2xHA-P2A-NLS-EGFP targeting vector was transfected into the abovementioned ESC lines along with pX330-puro^R^ or -blasticidin^R^ vector harboring guide RNAs targeting the *Otx2* C-terminus. 2 x 10^5^ ESCs were cultured overnight and transfected with 3μg of each plasmid using TransIT-293 (Mirus Bio 2700). Cells were replated at low density onto 10cm dishes with the corresponding antibiotic selection for 24-72 hours. Individual clones were screened by PCR followed by TOPO cloning (Invitrogen) and sequencing.

### Mouse ESC culture and epiblast conversion

ESCs were cultured in KO DMEM (Gibco 10829018) supplemented with 15% FBS (Gemini Benchmark), 2mM Glutamax (Gibco 35050079), 0.1mM nonessential amino acids (Gibco 11140076), 0.1mM 2-mercaptoethanol (Gibco 21985023), 1000U/ml leukemia inhibitory factor (LIF; in house), and 100μg/ml penicillin/streptomycin (Gibco 15140163) and maintained on a feeder layer of mitomycin C-treated mouse embryonic fibroblasts (MEFs) on gelatin-coated plates. For ESC to EpiLC conversion, ESCs were lifted using collagenase IV (Thermo Fisher 17104019), centrifuged for 4min at 120g, and dissociated with Accutase (Thermo Fisher 00455556). 15,000 cells/cm^2^ were seeded onto fibronectin (Millipore FC010; 16.7ug/ml)-coated plates in N2B27 media with 12.5ng/ml heat stable bFGF (Thermo PHG0360) and 20ng/ml Activin A (Peprotech 120-14P) (FA media) supplemented with 1% Knockout Serum Replacement (Gibco 10828010). EpiLCs were harvested or converted to EpiSCs after 48hrs differentiation. For EpiSC conversion EpiLCs were dissociated with TrypLE (Thermo Fisher 12605010) and plated onto gelatin-coated plates with feeders at 15,000 cells/cm^2^ in EpiSC media (FA supplemented with 175nM NVP-TNKS656 (Selleck S7238).

### Mouse definitive endoderm specification and foregut conversion

EpiSCs cultured between passages 2-7 were differentiated first to primitive streak/ endomesodermal progenitors (PS) and then to definitive endoderm (DE) as described (Medina-Cano et al., 2022). Briefly, EpiSCs were seeded onto 96, 48, or 24well plates coated with 10ug/ml Laminin 521 (StemCell Technologies 77003) in plating media. Plating media (chemically defined media, CDM) is 1:1 IMDM (Gibco 12440053) and DMEM/F12 with Glutamax (Gibco 10565018), 1% chemically defined lipid concentrate (Thermo Fisher 11905031), monothioglycerol (Sigma M6145), Apotransferrin (R&D 3188AT), 0.7 μg/ml insulin (Millipore

Sigma I0516), and polyvinyl alcohol (Sigma P8136) supplemented with 12.5ng/ml bFGF, 20ng/ml activin A, 175nM NVP, 1% KSR and 2 μM thiazovivin (Millipore Sigma SML1045). PS was induced 5-6 hours after seeding with CDM + 40ng/ml activin A and 3μM CHIR99021 (Biovision 16775). DE was induced 16 hours after PS with CDM + 100ng/ml activin A and 100nM LDN193189 (Reprocell 04-0074). Cells were washed 1x in PBS -/- between all media changes. To convert DE to antral gastric progenitors (posterior foregut), DE cultured for 48 hours was grown for an additional 48 hours in CDM with 100nM LDN193189 and 2% FBS. 2μM retinoic acid was added, and cells cultured for 24 hours before fixation and staining. Cells were treated with 500nM dTAG13 (Tocris 6605) in DMSO or vehicle control in culture media at the indicated amounts of time.

### Micropatterns

ESCs were converted to EpiLCs for 48hrs and seeded onto micropattern chips (CYTOO Arena A) as described (Morgani and Hadjantonakis, 2021). Briefly, 2-6 x 10^6^ EpiLCs were plated in EpiLC media onto micropattern chips coated with 10μg/ml Laminin 521. Media was replaced with FA after 2h and cells allowed to aggregate on micropatterns for 24hrs. 200 ng/ml Wnt3a (R&D 5036-WNP) was added to induce anterior differentiation. Chips were fixed and stained 48hrs after Wnt3a addition.

### Human ESC culture and genetic engineering DE conversion

Human MEL-1 and H9 ESCs were cultured on tissue culture plates coated with vitronectin (5 μg/ml) in Essential 8 Flex media with supplements and normocin (50 μg/ml) (“E8”). Cells were passaged with home-made EDTA Dissociation Solution (0.5 mM EDTA and 35 mM NaCl in PBS) during maintenance or with Accutase for freezing or when preparing single cell suspensions. Cells were frozen in Stem Cell Banker. After thawing, Y-27632 was added at 10 μM to the culture media for 24 hours. A targeting vector for generation of the human OTX2-dTAG allele was built by combining a 1252bp PCR amplicon (chr14:56,801,072-56,802,323 in hg38) generated from MEL-1 genomic DNA with a FKBPV-2xHA-P2A-NLS-EGFP cassette, using Gibson assembly. The resulting vector was introduced into MEL-1 and H9 hESCs together with the pX330 Cas9 vector expression a guide RNA (5’-GTACAGGTCTTCACAAAACC-3’) targeting the *OTX2* STOP codon using lipofection in mTeSR media mixed with CloneR solution (1:10). Transfected cells were selected for 48h with G418 (500 μg/ml), using mTeSR:CloneR (1:10). After selection, cells were seeded at clonal density and grown in E8 supplemented with CloneR (1:10) for three days and another five days in E8 without CloneR when colonies with undifferentiated morphology and homogenous nuclear EGFP fluorescence were picked. Clonal lines were expanded and characterized by flow cytometry (intensity and homogeneity of EGFP fluorescence), genotyping PCR (using the primers 5’-GGCAGGGGAAATTGTGTGTT-3’ and 5’-TCGTAAAGTTTCAGTGCAGCT-3’ which localize on either side outside of the *OTX2* homology region included in the targeting vector), by Sanger sequencing of the PCR amplicon and by IF using antibodies against HA and OTX2. Confirmed homozygous lines were karyotyped before use in differentiation experiments.

### Human DE specification

Single cell suspensions of hESCs were seeded in E8 supplemented with 10 μM Y-27632 onto cell culture vessel that were coated with Laminin-521 at 4°C for 24h to achieve a seeding density of 30-40%. The next day, cells were washed with prewarmed F12 based media and cultured for 24h in PS induction media Cells were then washed with prewarmed F12 based media and cultured for another 40h to 48h in DE induction media, with one media change after 24h of culture. Two protocols slightly differing in their use of signaling modulators were used. Protocol 1 uses the same PS and DE induction media applied to mouse DE. Protocol 2 uses PS induction media with 100 ng/ml Activin A, 2 μM CHIR99021 and 100 mM PIK90 (Selleckchem S1187) and DE induction media with 250 nM LDN1931189 (see also Fig. S5C). Both protocols use identical base media and overall experimental timing.

### Immunofluorescence

For micropatterns, chips were washed 2x in PBS -/-, fixed in 4% PFA for 15min, and washed again 2x in PBS -/-. Chips were stained as described. Primary antibodies used were rat anti-SOX2 (), goat anti-SOX17 (), and rabbit anti-FOXA2 (Abcam), all at 1:300. Secondary antibodies (Alexa Fluor, Life Technologies, and Jackson Immunoresearch) were used at 1:500. Micropattern chips were mounted on microscope slides with Fluoromount G (Thermo Fisher 00495802) and imaged on an LSM880 confocal microscope (Zeiss). For EpiSCs, DE and foregut, cells were washed and fixed in wells as above, blocked in PBS +/+ with 1% BSA, 0.1% Triton-X-100, and 3% donkey serum (Sigma D9663) and stained in primary antibodies overnight. Cells were washed 3x in PBS +/+ with 0.1% Triton X-100 (PBS-T) and then incubated in secondary antibodies for 1 hour. Cells were then again washed 3x in PBS-T. Hoechst33342 (Thermo Fisher H3570) was added to the last wash for 15 minutes for nuclear staining. Images were taken on an EVOS Cell Imaging System (Thermo Fisher) or Leica DMi8 microscope. Primary antibodies used were anti-SOX17 (R&D AF1924), anti-FOXA2 (Abcam 108422), anti-T (R&D AF2085), anti-HA (Abcam 9110) anti-OCT4 (Santa Cruz 8628), anti-GFP (Aves GFP1020), anti-SOX2 (Invitrogen 14981180), anti-PDX1 (Abcam 47308). For both micropatterns and directed differentiation, primary antibodies were used at 1:300 except anti-HA and anti-GFP (1:1000) and anti-PDX1 (1:100). Secondary antibodies (Alexa Fluor, Life Technologies, and Jackson Immunoresearch) were used at 1:500.

### Flow cytometry

Cells were washed with PBS -/-, dissociated with Accutase, resuspended in FACS buffer (80% PBS -/-, 20% FBS, 0.1% 0.5M EDTA), and incubated with indicated antibodies for 15min at room temperature in the dark. Cells were washed once in PBS -/-, resuspended in FACS buffer with DAPI, and analyzed on a BD FACS Canto. Data were analyzed using FlowJo. CXCR4 antibody conjugated to APC (eBioscience 17999182) and PDGFRa antibody conjugated to PE-Cy7 (eBioscience 25140180) were used at 1:200 dilution.

### RT-qPCR

Total RNA was extracted using TRIzol (Life Technologies 15596018) and purified with the RNA Clean and Concentrator kit (Zymo R1014). 1μg of RNA/sample was used for reverse transcription using the iScript kit (BioRad 1708841). qPCR was performed on cDNA samples in triplicate using PowerUp SYBR green PCR master mix (Thermo Fisher A25778) on an Applied Biosystems QuantStudio3. Primers used are in **Table S5**.

### Western blotting

Cells were washed with PBS -/- and harvested with Trypsin (Life Technologies 25200114). Nuclear lysates were prepared using the NE-PER kit (Thermo Fisher 78835) according to the manufacturer’s instructions. Western blots were performed using the following antibodies: anti-HA (Abcam 9110), anti-OTX2 (Abcam 21990), anti-histone H3 (Abcam 1791). Protein concentration was measured using Bradford Reagent (BioRad 5000201) and samples were boiled, resuspended in Laemmli Sample Buffer (BioRad), and run using BioRad PROTEAN System or Invitrogen precast gels.

### scRNA-seq

Cells at indicated stages were dissociated using Accutase to single-cell suspension in PBS-/-with 0.04% BSA. Samples were labeled with Cell Multiplexing Oligos (10x Genomics Chromium Next GEM Single Cell 3ʹ Reagent Kit v3.1 Dual Index) and live cells sorted using DAPI on a BD Influx Cell Sorter. Samples were then pooled for multiplexing in PBS-/-with 0.04% BSA (targeting 10,000 cells/sample) and single-cell libraries generated according to the manufacturer’s instructions (10x Genomics Chromium Single Cell Gene Expression - Cell Multiplexing). All cell sorts and deep sequencing were performed at the Weill Cornell Flow Cytometry and Weill Cornell Genomics Resources Core Facilities respectively.

### CUT&RUN

CUT&RUN on OTX2-HA was performed on 500,000 EpiSCs or DE cells, similar as previously described (Skene et al., 2018). For DE, live cells were selected by DAPI sort on an Influx Cell Sorter before performing CUT&RUN. BioMag Plus Concanavalin A beads (ConA beads, PolySciences NC1358578) were washed twice in 100μl cold Bead Activation Buffer (20mM HEPES pH7.5, 10mM KCl, 1mM CaCl_2_, 1mM MnCl_2_). 20μg of ConA beads per CUT&RUN reaction were used. Sorted cells were washed twice in Wash Buffer (20mM HEPES pH7.5, 150mM NaCl, 0.5mM Spermidine (Acros AC132740050), ½ Protease Inhibitor tablet) and bound to activated ConA beads in Wash Buffer for 10 minutes at room temperature. Samples were incubated in PCR strip tubes with 1:100 anti-HA (CST C29F4 #3724) or anti-IgG (EpiCypher 130042) in Antibody Buffer (Wash Buffer with 2mM EDTA and 0.01% Digitonin (Millipore Sigma 300410)) overnight with gentle rocking at 4°C. Samples were washed twice in Digitonin Buffer (Washer Buffer with 0.01% Digitonin) and ProteinA/G MNase (EpiCypher) was bound to samples for 1 hour at 4°C. ProteinA/G MNase was diluted at 20x in 50μl Antibody Buffer according to manufacturer’s instructions. Samples were washed twice in Digitonin Buffer again and MNase digestion was activated with 1μl 100mM CaCl_2_ in 50μl Digitonin Buffer for 2 hours. To extract bound chromatin fragments digestion was quenched using 33μl STOP Buffer (340mM NaCl, 20mM EDTA, 4mM EGTA, 50μg/ml RNAse A, 50μg/ml glycogen, 0.015 ng/μl E. coli spike-in) and samples incubated at 37°C for 20 minutes. Chromatin was separated from ConA beads by centrifuging 5 minutes at 16,000g and saving supernatant. DNA was purified by adding 0.1% SDS and 5μg Proteinase K for 10 min at 70°C, performing phenol-chloroform extraction, and precipitation in 100% ethanol at −80°C overnight. Pellets were then washed in 100% ethanol and resuspended in 12μl nuclease free water. CUT&RUN libraries were prepared using the Thruplex Library Prep kit (Takara) according to the manufacturer’s instructions. The number of cycles for library PCR amplification was determined by running 10% of each library sample in an Applied Biosystems qPCR machine with 0.25μl Eva Green dye (Biotium 31000) for 40 cycles and adding cycles to each sample as described (Buenrostro et al., 2015). Libraries were size selected with 1.5x volume SPRI beads (Beckman Coulter B23317) and sequenced at the Weill Cornell Genomics Resources Core Facility on an Illumina NovaSeq 6000 (PE-50, 30 million reads per sample).

### ATAC-seq

DE cultured with dTAG13 or DMSO vehicle for 24 hours was sorted for live cells on Influx as described for CUT&RUN. ATAC-seq was performed on three technical replicates of 50,000 cells each for DMSO or dTAG13-treated cultures as described (Murphy et al., 2023) using the Nextera DNA library preparation kit (Illumina FC121103) and NEBNext High-Fidelity 2X PCR Master Mix (NEB M0541). Libraries were sequenced on an Illumina NovaSeq 6000 (PE-50, 20 million reads per sample).

### Data analysis scRNA-seq analysis

Fastq files were processed using CellRanger (10x genomics cloud). The raw h5 file was further processed using CellBender to eliminate background reads, empty beads, and other artifacts (Fleming et al., 2022). Scanpy (Wolf et al., 2018) was used for downstream analyses. Only cells efficiently labeled with the demultiplexing primers were kept. Resulting cell clusters with high percentage of mitochondrial reads or a relatively low number of reads were eliminated. Palantir (Setty et al., 2019) was used for pseudo-time analyses. For differential expression analyses, Wilcoxon testing was performed and only genes with padj<0.05 and log2 fold-change>1 were considered as differentially expressed genes (DEGs). The scores/ranking per gene given by the scanpy.get.rank_genes_groups_df function were used.

### CUT&RUN sequencing analysis

CUT&RUN fastq files were processed as previously described (Uyehara et al., 2022) using a custom snakemake (v6.6.1) pipeline (Molder et al., 2021). Briefly, reads were trimmed using bbmap (v38.95) with parameters ktrim=r ref=adapters rcomp=t tpe=t tbo=t hdist=1 mink=11. Trimmed reads were aligned to the mm10 reference genome using Bowtie2 (v2.3.5.1) with parameters--local--very-sensitive-local--no-mixed--no-discordant--phred33-I 5-X 999 (Langmead and Salzberg, 2012). Reads with a MAPQ score less than 5 were removed with samtools (v1.14) (Li et al., 2009). PCR duplicates were removed with Picard (v2.26.10) using parameter VALIDATION_STRINGENCY=LENIENT. Fragments between 20 and 120bp were isolated using a custom awk script and used for downstream analyses as previously recommended (Skene and Henikoff, 2017). Bigwigs were generated from bam files using bedtools (v2.30) normalizing to sequencing depth using the formula: (effective_genome_size / read_count * read_length) with 1870000000 as the effective genome size (Quinlan and Hall, 2010). Bam files were converted to bed files with bedtools (v2.30) with parameter-bedpe. MACS (v2.2.6) was used to call peaks on individual and merged replicates using parameters-g 1870000000--nomodel--seed 123 –keep-dup all (Zhang et al., 2008).

### ATAC-seq

The raw ATAC-seq reads were trimmed using fastp to remove low-quality bases from reads (quality<20) and adapter sequences. The trimmed reads were aligned using Bowtie2 (Langmead and Salzberg, 2012) to UCSC genome assembly mm10. Duplicate reads were removed using Sambamba. Peaks were identified with MACS2 (Gaspar, 2018) and those overlapping with satellite repeat regions were discarded. For further analyses, a union peak atlas was created from the MACS2 files. Peak intensity for each sample was counted using featureCounts (Liao et al., 2014). HOMER (v4.11) (Heinz et al., 2010) was used for motif analyses. To associate peaks with nearby genes and genomic location, ChipSeeker (Yu et al., 2015) was used. To compare peaks from CUT&RUN and ATAC-seq experiments, *bedtools intersect* (Quinlan and Hall, 2010) was used with default parameters. The dataset published by (Medina-Cano et al., 2022) was used as a reference EpiSC dataset.

### Quantification and statistical analyses

Statistical analysis of flow cytometry and IF data was done in PRISM 9 (GraphPad), with specific tests and corrections applied indicated in the respective figure legends.

### Image analysis

Immunofluorescence images were analyzed using Fiji/ImageJ (Schindelin et al., 2012). For directed differentiations, fluorescence intensity was quantified using CellProfiler 4.2.1 (Carpenter et al., 2006). Cells were identified by DAPI staining using the IdentifyPrimayObjects module and mean fluorescence intensity in each stained channel was measured using the MeasureObjectIntensity module. Statistical significance was calculated in PRISM 9 (GraphPad) using one-way ANOVA. For micropatterns, entire chips were imaged using a 5x objective and 11×11 tiling. Individual colonies were detected based on DAPI, size and circularity thresholds using custom Fiji macros. The mean pixel intensity for each fluorescent channel for each colony as well as the mean and standard error across all colonies was calculated in MATLAB R2021a (MathWorks) and normalized by the maximum value from each channel using custom scripts (available upon request). For quantification of posterior foregut progenitors, SOX2^+^PDX1^+^ cell clusters on 24-or 48 well cell culture plates were imaged and counted at 10X magnification on a Leica DMi8 microscope. Cell clusters were quantified for 5 replicates per condition (dTAG-13 added at PS or after 24 hours DE differentiation; and DMSO control). Cluster counts for each replicate were divided by the respective growth area (cm^2^) to normalize for plate well area over two independent experiments. Significance was calculated using the Kruskal-Wallis nonparametric test.

---

**Figure S1.**
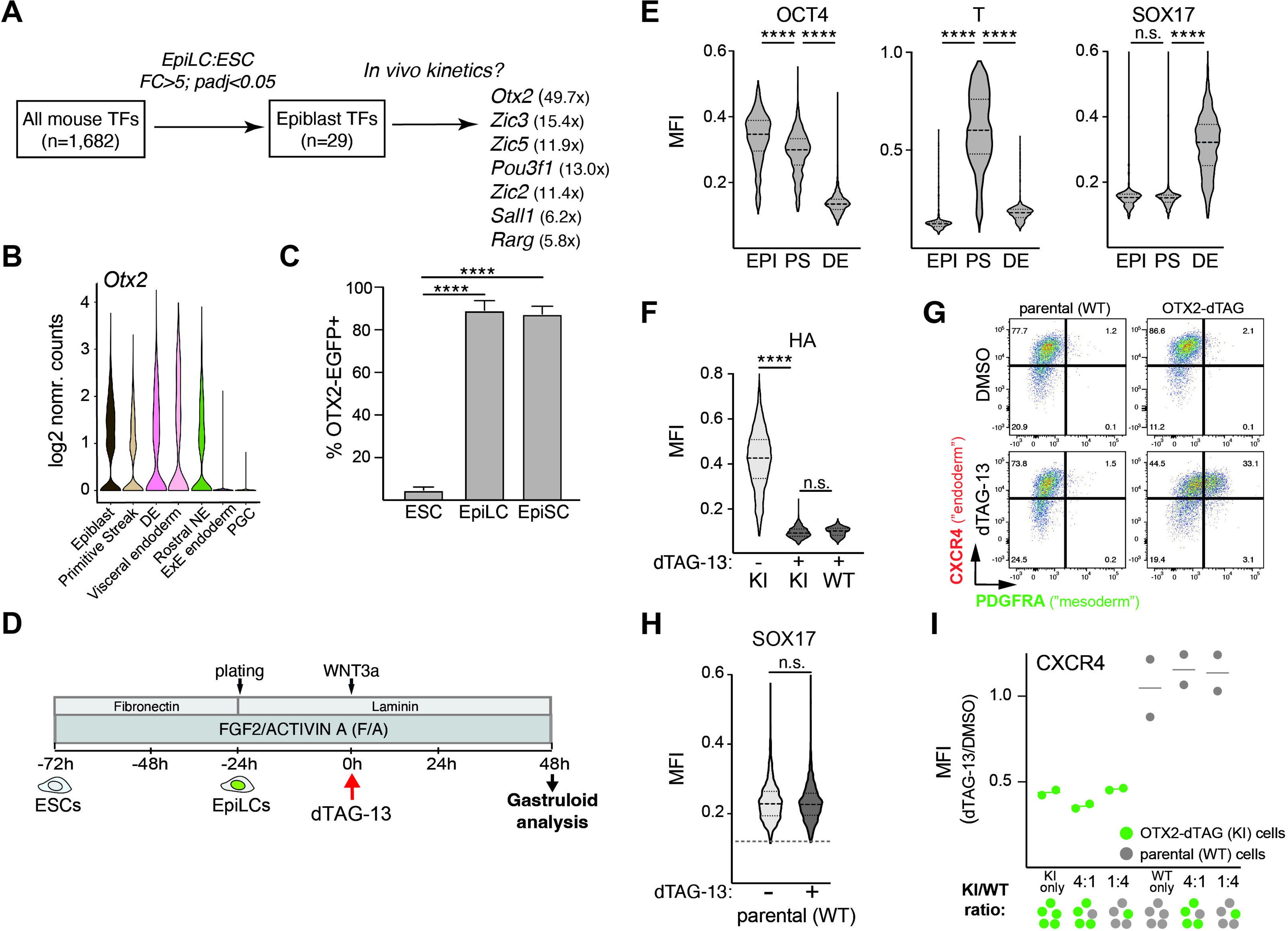
OTX2 depletion impairs DE development in murine models of germ layer specification. A. *In silico* strategy to identify candidate transcriptional regulators of germ layer specification from a recently assembled list of 1,682 TFs encoded in the mouse genome (Garipler et al., 2022). B. *Otx2* expression levels in DE and in tissues with known *Otx2* functions (visceral endoderm, rostral neuroectoderm) present in gastrulation-stage mouse embryos (Pijuan-Sala et al., 2019). Extra-embryonic endoderm (ExE endoderm) and PGCs are shown as representative tissues not expressing *Otx2*. C. Percentage Otx2-EGFP^+^ cells in cultures of mESCs, EpiLCs and EpiSCs as measured by flow cytometry. (**) p<0.01 with two-way ANOVA with Tukey’s multiple comparison test (n=3 independent cultures). D. Experimental strategy for the generation of anterior micropatterns (“gastruloids”) with timing of dTAG-13 treatment indicated. E. IF quantification of intensity of stage-specific marker proteins at indicated stages of differentiation. (****) p<0.0001 with one-way ANOVA with Tukey’s multiple comparison test. N > 500 nuclei were analyzed for each marker and sample. F. Quantification of OTX2 (via HA IF) in PS cultures after 1h of exposure to dTAG-13 or DMSO. (****) p<0.0001 with one-way ANOVA with Tukey’s multiple comparison test. N > 500 nuclei were analyzed for each marker and sample. G. Representative flow cytometry plots showing CXCR4 (endoderm marker) and PDGFRA (mesoderm marker) expression in DE established from OTX2-dTAG EpiSCs and EpiSCs derived from parental wildtype (WT) mESCs in presence of DMSO (top panels) or dTAG-13 (bottom panels). H. IF quantification of intensity of SOX17 protein in DE established from WT EpiSCs cultured in presence of DMSO or dTAG-13. Statistics with unpaired t-test. I. Ratio of CXCR4 levels in presence of DMSO or dTAG-13 as measured by flow cytometry on OTX2-dTAG (KI) and WT DE in mixed cultures of the indicated ratios. OTX2-dTAG were distinguished from WT cells based on EGFP expression.

**Figure S2.**
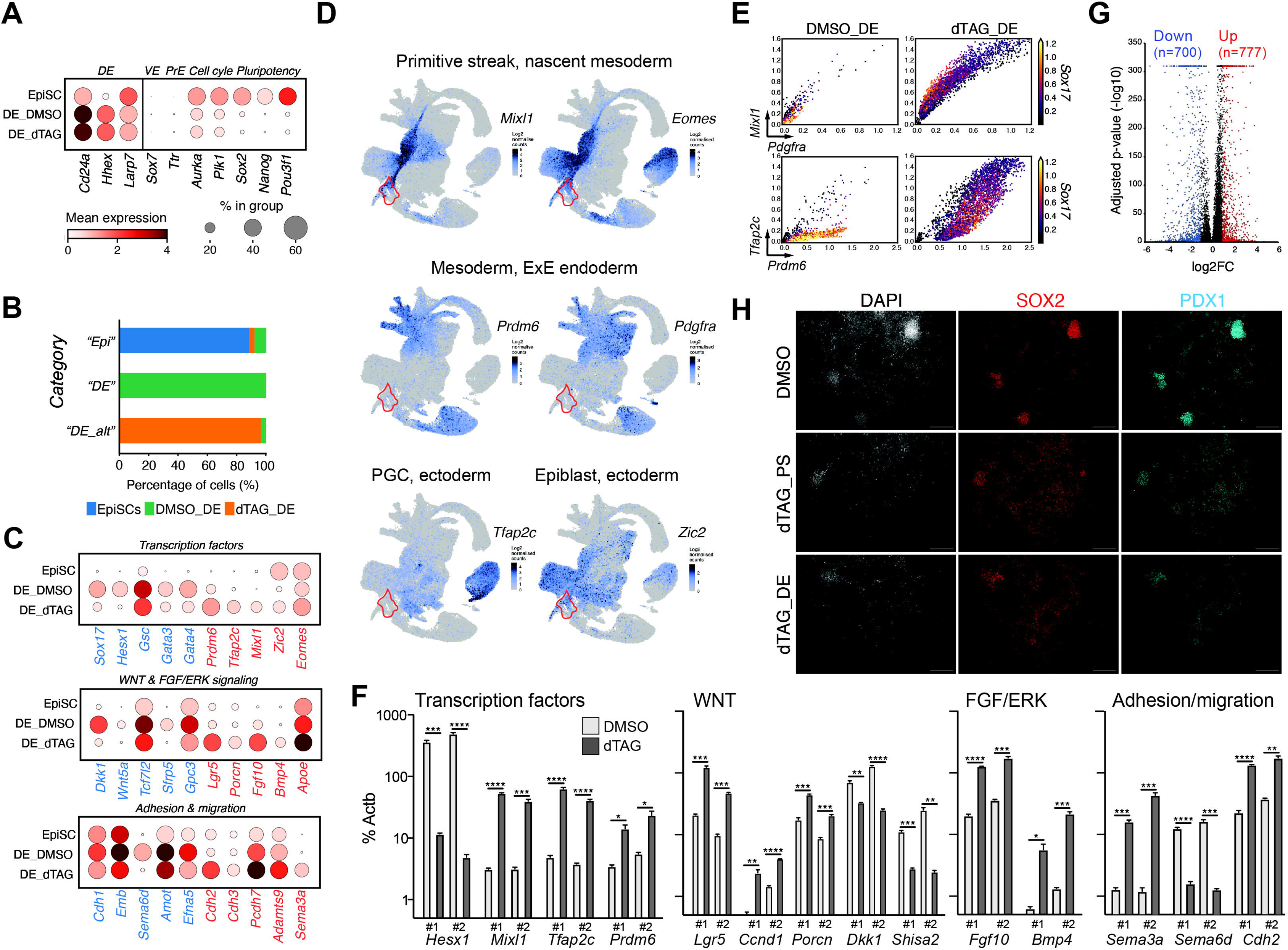
Figure 2. Transcriptional dysregulation in DE upon OTX2 depletion. A. Expression levels of select genes not affected by OTX2 depletion, including a subset of DE markers (*CD24a*, *Hhex*, *Larp7*) as well as markers for visceral endoderm (*Sox7*), primitive endoderm (*Ttr*), cell cycle progression (*Aurka*, *Plk1*) and primed pluripotency (*Sox2*, *Nanog*, *Pou3f1*). B. Abundance of cells assigned an “Epi”, “DE” or altered DE (“DE_alt”) identity in indicated cultures based on scRNA-seq analysis. C. Expression levels of select DEG^DOWN^ (blue) and DEG^UP^ (red) after OTX2 depletion at PS, representing developmental TFs, signaling regulators and proteins involved in cellular adhesion or migration. D. Expression in gastrulation-stage embryos (from Pijuan-Sala et a., 2019) of select TFs associated with indicated non-DE lineages that are upregulated in DE upon OTX2 depletion. The red outline indicates position of DE in the embryonic UMAP. E. Correlation between expression levels of *Sox17* (DE marker) and levels of indicated non-DE markers in DE cultures established in presence of DMSO (left panels) or dTAG-13 (right panels). F. Validation by qPCR of the effect of OTX2 depletion on select DEG^DOWN^ (blue) and DEG^UP^ (red) in two independent cell lines. (*) p<0.05, (**) p<0.01, (***) p<0.001 and (****) p<0.0001 with multiple t-tests and Bonferroni-Dunn correction. N = 3 measurements. G. Volcano plot showing expression changes in OTX2^depl_DE^ DE with DEGs (logFC>1;padj<0.05) in blue (DOWN) or red (RED), respectively. H. Representative IF images after staining of gastral foregut cultures exposed to DMSO or dTAG-13 at PS or DE with SOX2 and PDX1 antibodies.

**Figure S3.**
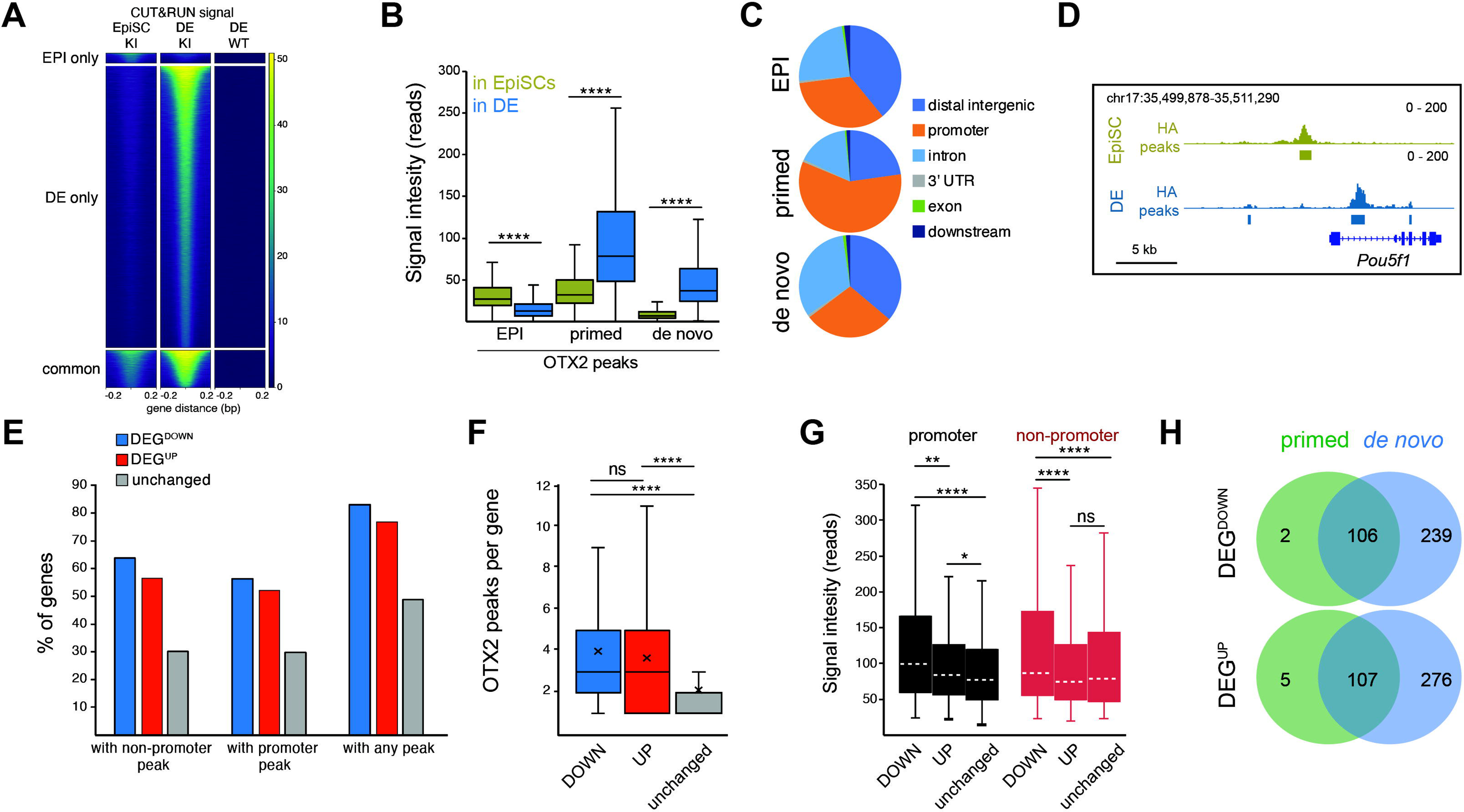
Dynamics of OTX2 genome occupancy during DE specification. A. Tornado plot showing signal intensity at HA CUT&RUN peaks called in OTX2-dTAG EpiSCs and DE as well as in corresponding genomic regions in WT (non-transgenic) DE. B. Cell-type specific (EpiSCs and DE) HA CUT&RUN intensity at EPI, primed and *de novo* OTX2 peaks. (****)p<0.0001 with paired Wilcoxon test. C. Genomic distribution of EPI, primed and *de novo* HA CUT&RUN peaks. D. IGV tracks showing cell type-specific OTX2 binding at the pluripotency-associated *Pou5f1* locus, a known target of OTX2 in primed pluripotent cells. E. Fraction of DEG^DOWN^, DEG^UP^ and control genes unaffected by OTX2 depletion that have an associated OTX2 peak in DE. F. Average number of OTX2 peaks (in DE) at OTX2 bound DEG^DOWN^ (n=327), DEG^UP^ (n=345) and unaffected gene loci (n=572). (****)p<0.0001 with Mann-Whitney test for unpaired data, two side and confidence interval of 95%. G. Intensity of HA CUT&RUN signal at promoter and non-promoter OTX2 peaks associated with DEG^DOWN^ (n=360 for promoters; n=942 for non-promoters), DEG^UP^ (n=344 for promoters; n=915 for non-promoters) and unaffected control genes (n=451 for promoters; n=752 for non-promoters). (*)p<0.05, (**)p<0.01 and (****)p<0.0001 with Mann-Whitney test for unpaired data, two side and confidence interval of 95%. H. Overlap of OTX2-bound DEG^DOWN^ (top) and DEG^UP^ (down) gene loci with primed OTX2 peaks (green) and such loci with *de novo* OTX2 peaks (blue).

**Figure S4.**
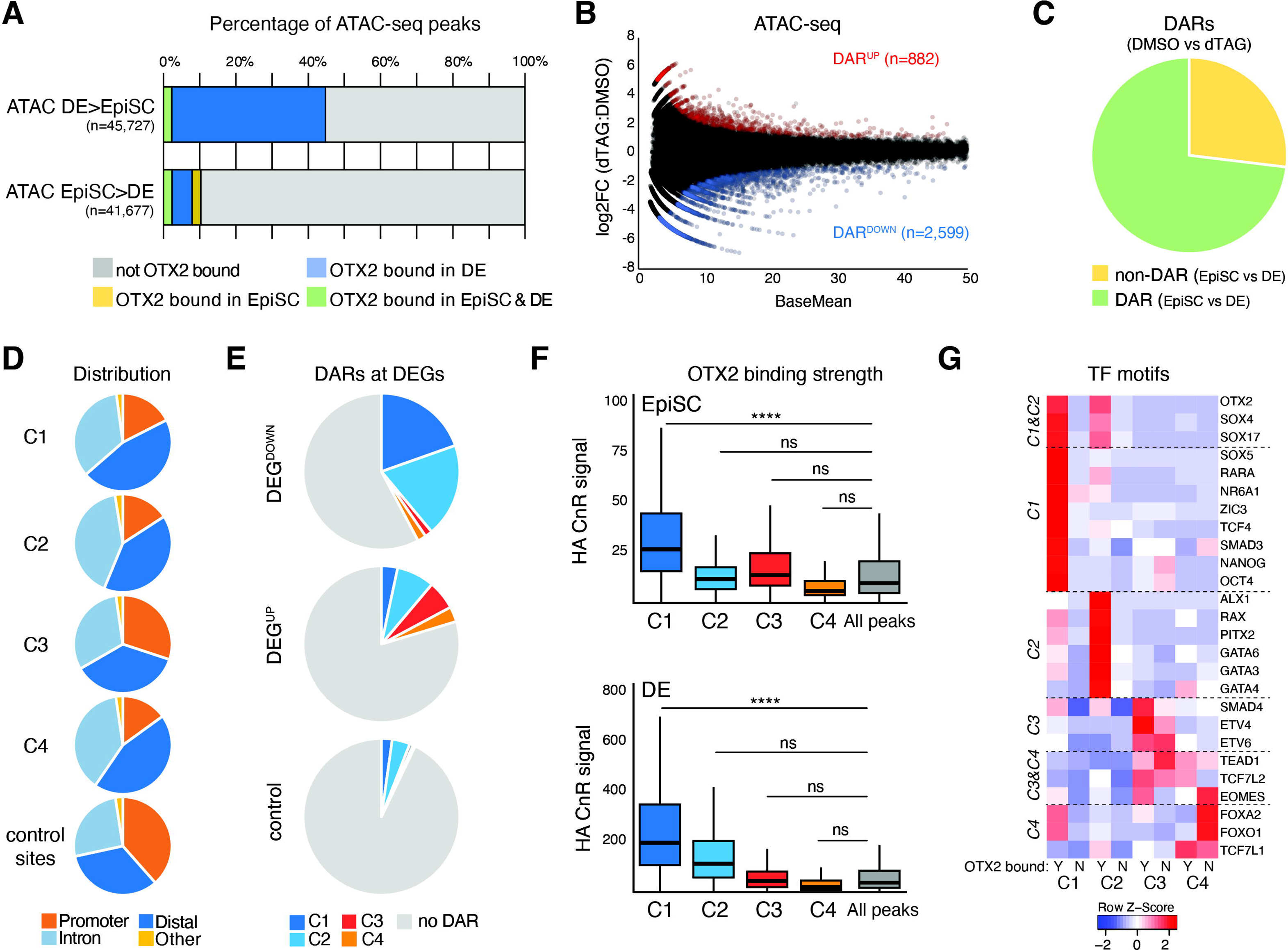
OTX2-dependent remodeling of chromatin accessibility during DE specification. A. Percentage of genomic regions with differential chromatin accessibility between EpiSCs (Medina-Cano et al., 2022) and DE that are bound by OTX2 in either, both or neither cell type. B. MA plot showing ATAC-seq signal change in DE derived in either presence of DMSO or dTAG-13. Significantly different (logFC>1;padj<0.05) DARs are indicated in red (UP) or blue (DOWN). C. Fraction of regions that change their chromatin accessibility in DE upon OTX2 depletion (DARs DMSO vs dTAG) that undergo accessibility changes during the transition from EpiSCs to DE (green) or not (yellow). D. Genomic distribution of genomics regions in DE that are affected in their chromatin accessibility by OTX2 depletion at PS (C1 to C4). E. Fraction of DEG^DOWN^, DEG^UP^ and unaffected control genes with an associated C1-C4 DAR or no associated DAR. F. Intensity of HA CUT&RUN signal at C1 to C4 DARs in EpiSCs (top) and DE (bottom). Wilcoxon test was used for statistics (each group versus the ensemble of peaks per cell type) with (****)p<0.0001. G. Supervised clustering of TF motif enrichment at C1-C4 DARs, distinguishing between DARs bound by OTX2 and DARs not bound by OTX2. Dotted lines indicate TFs exhibiting enrichment at similar DAR categories.

**Figure S5.**
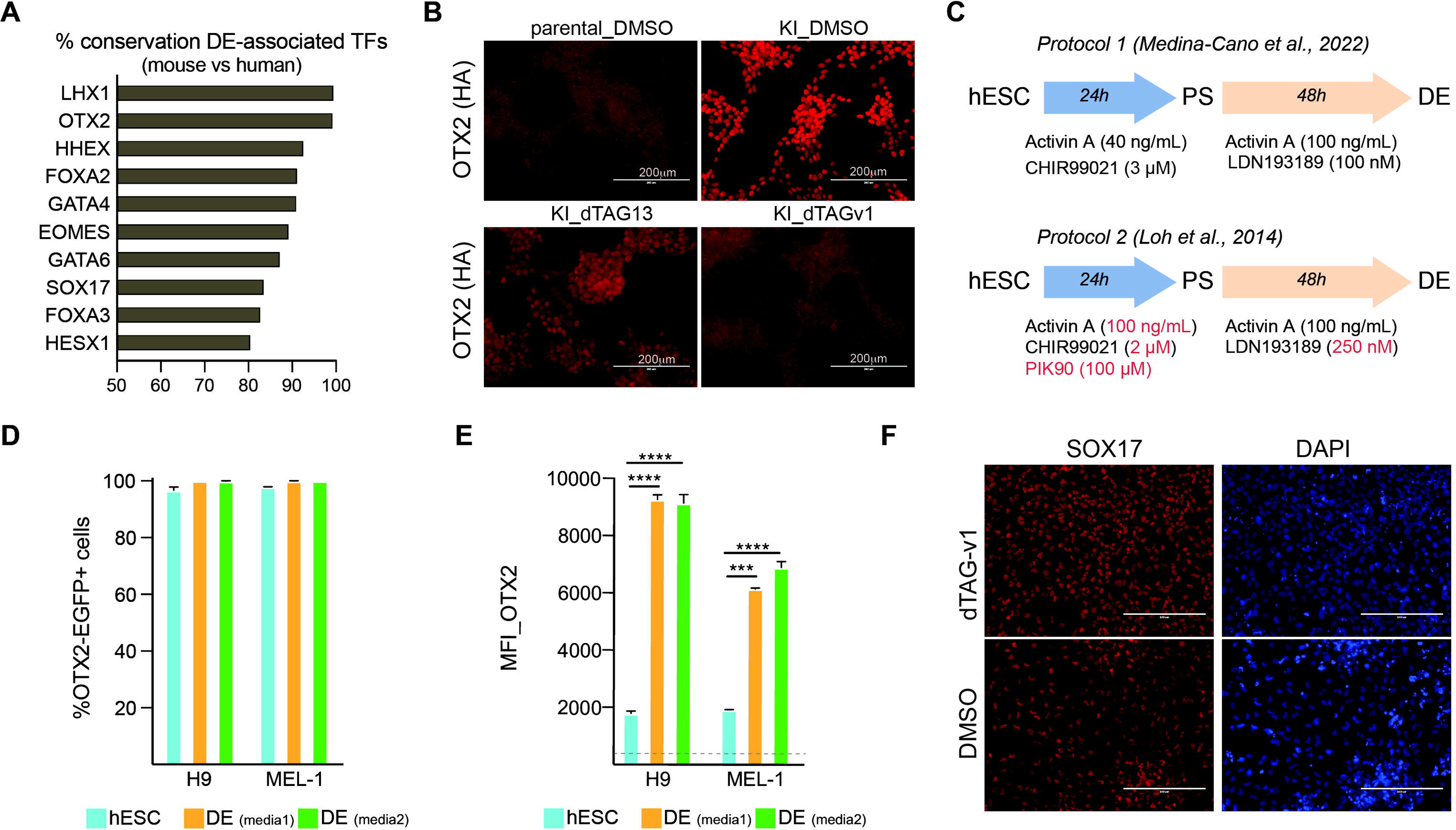
Impaired human DE formation upon OTX2 depletion. A. Select DE-associated TFs ranked by their degree of protein conservation between mouse and human. B. Anti-HA IF of OTX2-dTAG hESCs exposed to either DMSO, dTAG-v1 or dTAG-13 for 2h. Parental hESCs cultured in DMSO are shown to represent background fluorescence levels. Note residual retention of HA signal in presence of dTAG-13. C. Outline of the two differentiation regimens for the generation of DE from hESCs employed in this study. Differences in compounds used or concentrations applied are highlighted in red. D. Percentage of OTX2-EGFP^+^ cells in OTX2-dTAG hESCs and derivative DE. E. OTX2-EGFP expression levels (as measured by flow cytometry) in hESCs an DE (derived in two different media compositions) of indicated backgrounds. The grey dotted line indicates background fluorescence levels measured in parental, non-transgenic hESCs. (***) p<0.001 and (****) p<0.0001 with one-way ANOVA with Tukey’s multiple comparison test. F. Representative IF images after staining human DE derived in either presence of dTAG-v1 or DMSO with anti-SOX17 antibody or DAPI.

